# Deciphering molecular mechanisms of synergistic growth reduction in kinase inhibitor combinations

**DOI:** 10.1101/2024.03.12.584561

**Authors:** Eirini Tsirvouli, Ana Martinez del Val, Liv Thommesen, Astrid Lægreid, Martin Kuiper, Jesper Olsen, Åsmund Flobak

**Affiliations:** Department of Biology, Norwegian University of Science and Technology, Trondheim, Norway; Department of Clinical and Molecular Medicine, Norwegian University of Science and Technology, Trondheim, Norway; The Novo Nordisk Foundation Center for Protein Research, University of Copenhagen Denmark, Copenhagen, Denmark; Department of Biomedical Laboratory Science, Norwegian University of Science and Technology, Trondheim, Norway; The Cancer Clinic, St Olav’s University Hospital, Trondheim, Norway; Department of Biotechnology and Nanomedicine, SINTEF Industry, Trondheim, Norway

## Abstract

In cancer treatment, the persistent challenge of unresponsiveness of certain patients to drugs or the development of resistance post-treatment remains a significant concern. Drug combinations that synergistically reduce tumor growth emerge as a promising avenue to address this issue. Here, we aimed to characterize the mechanism of action of two synergistic drug combinations that target PI3K together with MEK1 or with TAK1 and used time course measurements of phosphoproteomics and transcriptomics in response to single inhibitors and their combinations. Our analysis untangled those responses driven by single drugs and responses that were unique to the combinations. We observed a high overlap between single-drug responses and their combinations, suggesting that single-drug mechanisms dominate the mechanism of action of the combinations of the kinase inhibitors. Despite a high overlap, both drug combinations exhibited a synergistic modulation of several cell fate regulators found at the convergence points of the targeted pathways, including the key regulator of intrinsic apoptosis BCL2L11. Interestingly, the responses in both combinations were largely limited to the targeted pathways, namely PI3K/AKT and MAPKs, with very limited change of any other additional cell fate decision pathways. In addition, we observed a strong downregulation of nucleotide metabolism and tRNA biosynthesis uniquely in the combinations, which could be attributed to the reduced activity of mTOR and ATF4. Our approach provides insights into the molecular mechanisms affected by the PI3Ki-TAK1i and PI3Ki-MEKi combinations and can serve as a flexible framework for dissecting drug combination responses based on multi-omics measurements.

## Introduction

Personalized medicine and targeted drug therapies represent a paradigm shift in cancer treatment. These approaches provide the potential to tailor treatments specifically to the patient, resulting in improved treatment effectiveness and lowered incidence of adverse outcomes. Targeted therapies, usually based on small molecules or monoclonal antibodies, interfere with molecular entities that have an indispensable role in disease development. Whereas chemotherapy and radiotherapy impact both cancerous and noncancerous cells indiscriminately by affecting all actively growing cells, targeted therapies can be designed to specifically target cell fate signaling that is central to cancerous cell growth ^1^. Most small molecule-based therapies focus on inhibiting kinases that are abnormally active in cancers. The advent of kinase inhibitors has dramatically changed the field of oncology treatment ^2^, with currently some 72 kinase inhibitors approved by the FDA ^3^.

Insights into the mechanism of drug action are essential to understanding how drugs can best be used in combinations for new potential therapeutic applications ^4,5^. Moreover, many cancers can develop resistance to a drug treatment, which poses an additional challenge for effective therapies. Drug combination therapies and drug synergies can improve cancer treatment either by increasing efficacy, reducing side effects, or preventing treatment resistance ^6^. One example is T-cell acute lymphoblastic leukemia (T-ALL), where NOTCH1 is a key oncogenic driver ^7^. The development of small molecule therapies to prevent NOTCH1 activation, such as γ-secretase inhibitors (GSIs), faced significant problems due to the development of drug resistance. In-depth analysis of the development of such resistance using Mass Spectrometry (MS)-based (phospho)proteomics has identified other targets in T-ALL, such as protein kinase C (PKC) delta, suggesting that the PKC inhibitor sotrastaurin can improve the anti-leukemic activity of the GSIs when provided in combination ^8^. Such examples highlight that drug combinations can be an alternative when monotherapies fail to provide a persistent therapeutic response.

Comprehending the effects of drug combinations is a complex endeavor, as the individual drugs can work together either in an additive, synergistic, or even antagonistic manner. Synergistic interactions, where the combined effect surpasses the sum of individual drug effects ^9^, are particularly intriguing, as they offer the potential for amplified therapeutic benefits. There are several mechanisms through which drug synergies can occur, including but not limited to the modulation of signaling pathways involved in the development of the disease, increasing the sensitivity to drug responses, and potentially inhibiting drug resistance mechanisms. The systematic characterization of the responses to synergistic drug combinations and the analysis of their mode of action may provide insight into how synergies arise. Such knowledge could potentially be useful to enable the rational design of efficient drug synergy screening ^10^.

Despite the potential advantages of potent drug combinations, their discovery and effective application in clinical treatments pose formidable challenges. While automated high-throughput screening platforms can uncover combinatorial effects, drug combination screens remain a resource-intensive and time-consuming endeavor, demanding extensive dose-response data. To streamline this process, computational simulations, and machine learning algorithms can be used to enable the preselection of drug combinations in silico, identifying those most likely to yield beneficial effects. Such approaches can significantly reduce the number of combinations requiring experimental testing ^11^. Previously, we employed logical modeling of cancer signaling networks for drug synergy identification in gastroenterological cancers ^12–14^. Among 21 pairwise combinations of seven kinase inhibitors, we identified and experimentally validated four synergies in the AGS cell line ^12^. Notably, the dual inhibition of TAK1 together with PI3K or with AKT1, a previously unreported synergy, demonstrated its potential for reducing tumor growth in both in vitro and in vivo experiments. Additionally, we confirmed the synergistic effects of MEK inhibition in combination with either PI3K or AKT1, aligning with prior experimental findings ^15^. The PI3K/AKT and MAPK pathways are central regulators of oncogenesis and tumor maintenance, and the combined blockade of the two pathways has been shown to act synergistically on a variety of tumors and to abolish resistance mechanisms ^16–18^. However, the combinatorial targeting of TAK1 with either PI3K or AKT has not yet been studied in great detail at a mechanistic level.

In this work, we aimed to characterize the mechanisms of two synergistic combinations of kinase inhibitors, namely the inhibition of PI3K jointly with the inhibition of MEK or with the inhibition of TAK. We conducted a comprehensive analysis by measuring time-course responses in phosphoproteomics and transcriptomics for both single inhibitors and their combinations. Our investigation untangled the distinct responses induced by individual drugs from those unique to the combination treatments, shedding light on how the profiles of the single drugs interact to produce the synergistic reduction in cell growth and cell cycle arrest.

## Results

### A multi-omics characterization of responses to PI3K-MEK and PI3K-TAK combined inhibition

We investigated the molecular mechanisms behind the synergistic cell growth reduction when combining a PI3K inhibitor with either TAK1 or MEK inhibitors in AGS gastric adenocarcinoma cells, examining temporal responses at the phosphoproteomics and transcriptomics levels (Figure 1). AGS cells were treated with individual inhibitors: TAK1 inhibitor (TAKi, 0.50 µM), PI3K inhibitor (PI3Ki, 0.70 µM), and MEK inhibitor (MEKi, 0.035 µM), along with the PI3Ki-MEKi and PI3Ki-TAKi combinations (Figure 1). Each drug, at these concentrations, has been observed to reduce growth by 50% after 48 hours compared to vehicle dimethyl sulfoxide (DMSO) controls ^12^. Phosphorylation changes were monitored at 30 minutes, 2 hours, and 8 hours to detect early signaling events, with a proteome analysis at the same time points to rule out that changes in phosphorylations were observed due to changes in total protein (Figure 1B). RNA-Seq analysis at 1, 2, 4, 8, and 24 hours was performed to assess the effect of the inhibitors on gene expression (Figure 1B). Beginning with a high-level assessment, we explored the effects at the pathway and process levels through overrepresentation analyses. Subsequently, we continued into the effector level, estimating kinase and transcription factor activities to discern key regulatory elements. Finally, we narrowed our focus to identify individual entities downstream of these effectors, aiming to elucidate their roles in governing cell fate decisions. This systematic workflow ensures a thorough exploration, and the results will be presented in this sequential order.

**Figure 1.**
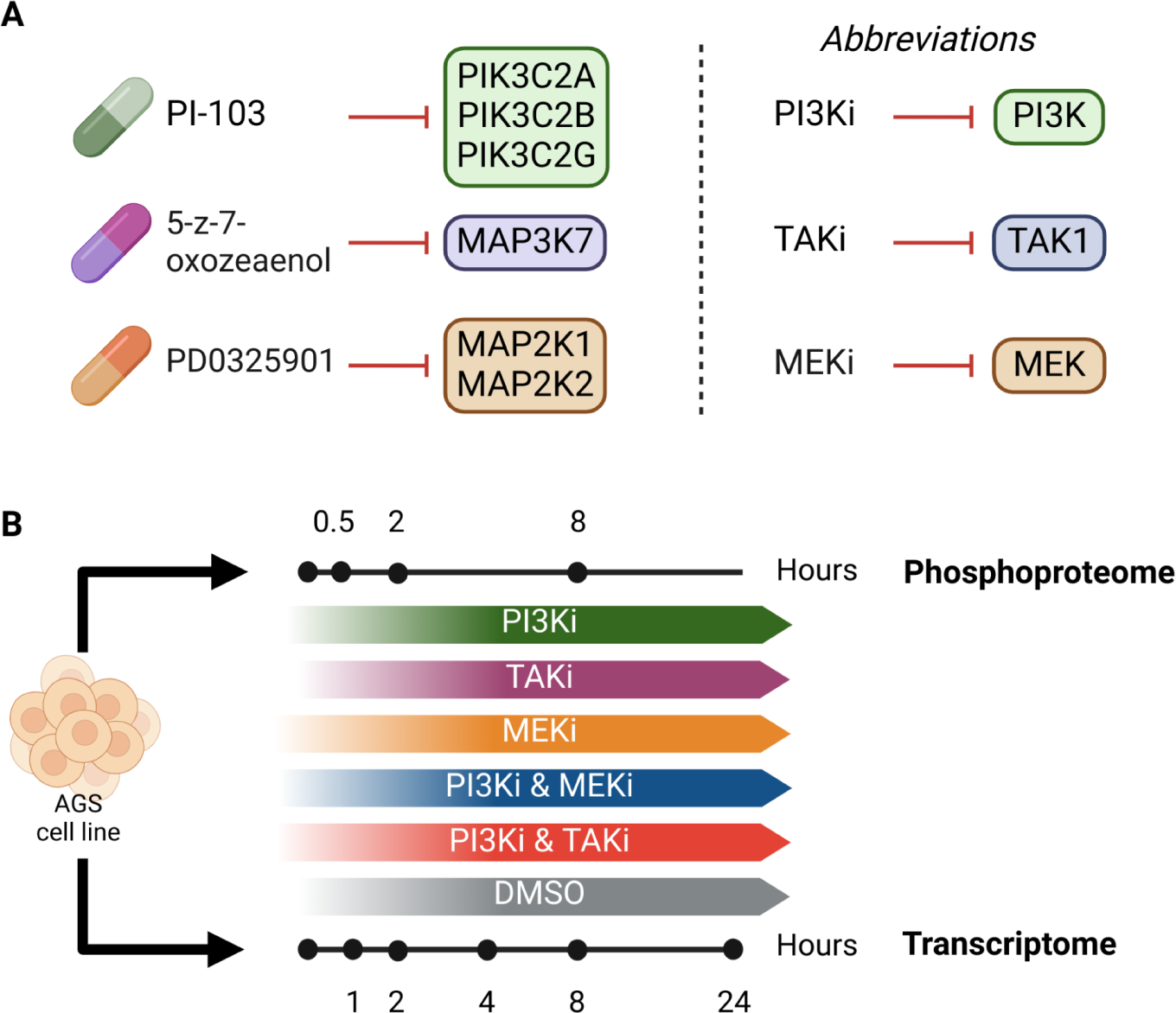
Overview of the experimental design. (A) Kinase inhibitors used in the study. In the left panel, the inhibitors are presented together with their main targets. In the right panel, the abbreviations used for each of the inhibitors and their targets throughout the manuscript are provided. (B) The gastric adenocarcinoma cell line (AGS) was used to investigate the effect of the combination of PI3Ki (inhibition with PI-103) with MEKi (inhibition with PD0325901) or with TAK1i (inhibition with 5Z-7-oxozeaenol.) Phosphoproteomics were measured at 0.5 hours, 2 hours, and 8 hours post-treatment. Transcriptomics were measured at 1, 2, 4, 8, and 24 hours post-treatment.

First, to identify early regulatory events and study the possible convergence and divergence in the signaling pathways triggered by each drug, or combination of drugs, we designed a mass spectrometry (MS) based phosphoproteomics analysis (Figure 2A). The drug effect at each time point was referenced to the corresponding effect of the vehicle at the same time point. Overall, we were able to quantify 18,517 phosphorylation sites, including 152 kinase activity regulatory sites ^19^ for 72 kinases. Principal Component Analysis (PCA) of the phosphoproteomes revealed a significant difference between DMSO and untreated at time zero samples, reflecting the importance of using DMSO-treated cells as a control in short-time phosphoproteomics signaling experiments (Figure 2B). Because of the depth of the phosphoproteomics profiling, the phosphorylation status of downstream targets of the inhibited kinases served as a positive control for the treatment effect. To begin with, the inhibition of MEK by PD0325901 is reflected by a reduction of ERK phosphorylation levels. As others have reported, we observed upstream MEK1/2 hyper-phosphorylation, which has been ascribed to the allosteric inhibitory nature of PD0325901, which in turn results in an interaction between upstream MEK1/s and c-RAF that triggers the hyperphosphorylation ^20^. This pattern of hyper-phosphorylation is recapitulated in our data, both in MEKi and in MEKi-PI3Ki combined treatment, where the regulatory tyrosine in position 187 of MAPK1 (ERK2) is rapidly down-regulated upon 30 minutes of drug treatment. Still, serine 226 in upstream MAP2K2 (MEK2) is upregulated consistently after 8 hours of treatment (Figure 2C). The 5Z-7-oxozeaenol inhibitor used against TAK1 has been found to have MEK1 as a secondary target but with a lower affinity compared to TAK1 ^21^. Interestingly, while TAK1 alone did not affect ERK1/2 activity (based on its phosphorylation status), the PI3Ki-TAKi combination induced the inhibition of the ERK2 activation site (Tyr-187) (Figure 2C). On the other hand, AKT1, which is a target of PI3K, shows inhibition on its regulatory sites in all treatments, although this effect seems to be less evident in the combined PI3Ki-MEKi treatment. Other downstream kinases of the PI3K-mediated pathway, such as mTOR, are only downregulated in the treatments containing PI3Ki. Finally, MAPK14 (p38), a downstream kinase of the TAK1 pathway, shows the greatest inhibition upon 8 hours of TAKi treatment, while in the combined PI3Ki-TAKi treatment, the effect, although smaller, appears already after 2 hours of treatment (Figure 2C). Overall, the observed changes agree with the expected changes based on the inhibitors’ target.

**Figure 2.**
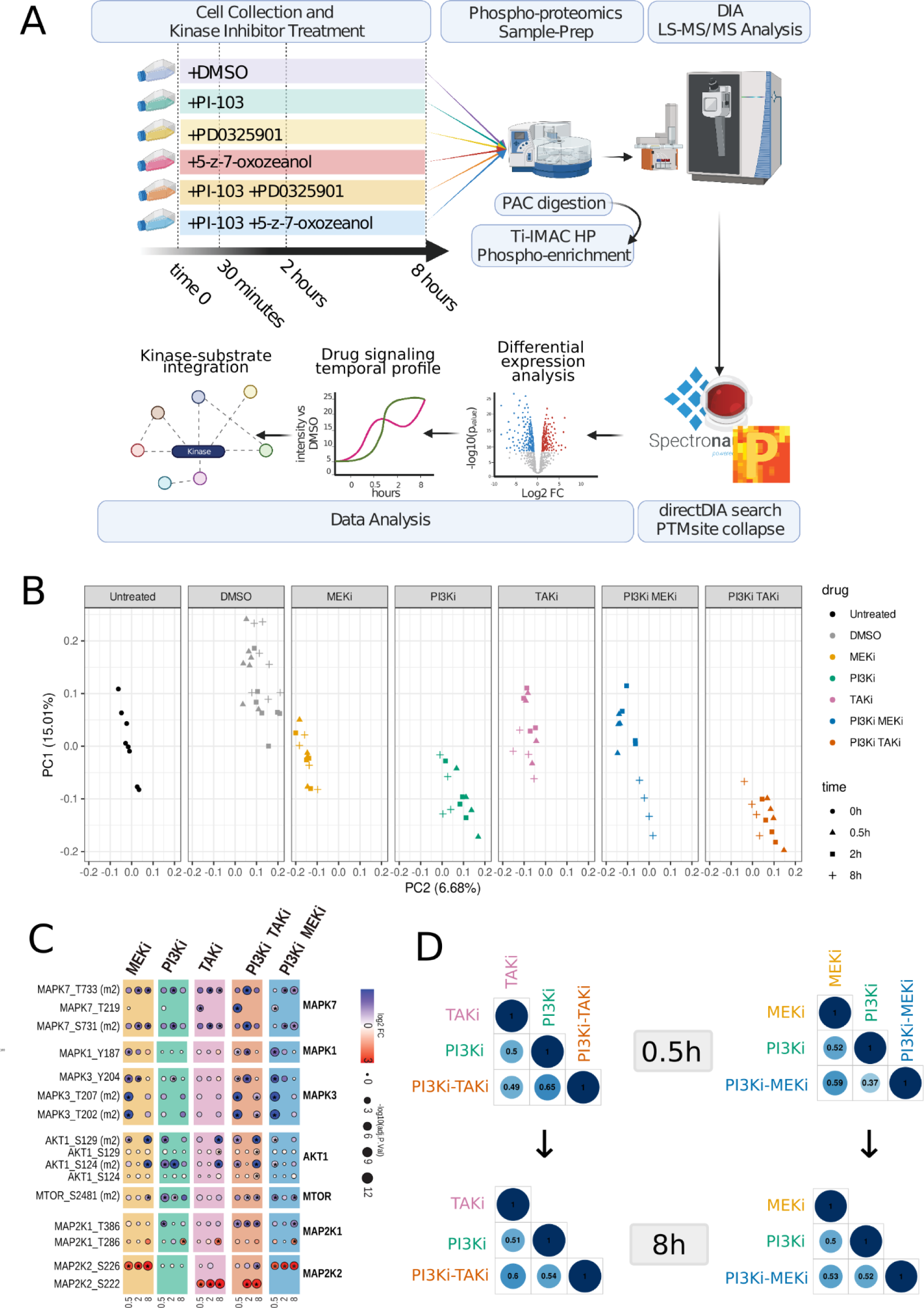
Phospho-proteomics analysis reveals the specific kinase targets of each drug and the convergence of drug synergies in kinase activation sites. (A) Experimental design for phosphoproteomics analysis. (B) Principal Component Analysis of phosphoproteome profiles of samples treated either with DMSO or with each drug combination and time zero (non-treated samples). (C) Most relevant kinase activation sites of the targeted pathways of each drug. The color of the dot indicates the log fold change of the intensity of the phosphorylation site in each drug treatment versus DMSO at the same time point. The size of the dot indicates the statistical significance of the change (two-sided, moderated two-sample t-test, FDR corrected by Benjamini-Hochberg). * indicates q-values < 0.05. (D) Pairwise correlation between the log fold change (logFC) between the single drugs and the combinations at 0.5 hours (upper plots) and 8 hours (bottom plots). The numbers within each circle show the correlation coefficient between logFCs.

Furthermore, we assessed how responses in single drug-treated cells compared to the responses in cells treated with their combinations. The logFC compared to DMSO for each phosphosite was plotted between individual drugs and synergies. All phosphorylation responses were positively correlated, with all correlation coefficients increasing with time (Figure 2D). Specifically, PI3Ki-MEKi had a higher correlation with MEKi (Pearson coefficient = 0.6), suggesting MEKi’s predominant role at early time points, with PI3Ki having a weaker positive correlation (0.4). Conversely, both PI3Ki and TAKi had a high contribution to PI3Ki-TAKi phosphorylation status, with Pearson coefficients of 0.65 and 0.5, respectively (Figure 2D). This suggests that the two combinations we have investigated arise through the mixed effect of additive and synergistic interactions between the two single drugs. Additionally, even though the PI3K inhibitor was common to both combinations, it appears that its contribution to the combination profiles varies depending on its partner.

### Temporal responses in phosphoproteomics reveal few differences in kinase activity between single drugs and combinations

Our data point to an overlap in the downstream signals regulated by the combined treatments, and temporal phosphoproteomics profiling can help figure out specificities in each treatment due to the kinetics or magnitude of the regulation. To explore the kinetics of each treatment and differentiate between additive and synergistic effects, we performed k-means-based clustering using the log2 fold changes at each time point of treatment compared to vehicle control (Figure 3). For each cluster of a combined treatment (separately for PI3Ki-TAKi and PI3Ki-MEKi), we also provide the plots of the corresponding values of the phosphosites observed in the individual treatments (Fig. 3A, left part of individual panels). Most of the temporal kinetics observed in the combined treatments are also found in treatments with one or sometimes even both of the single inhibitors. These results suggest that many of the effects observed in the combined treatments likely reflect the cumulative impact of the two drugs. However, some clusters, such as cluster #9 in PI3Ki-TAKi and PI3Ki-MEKi, show a transient effect in the combined treatments, while with each individual drug, the effect is longer lasting (Figure 3A). Another interesting difference between the two different combination treatments can be observed upon kinase motif annotation analysis (Figure 2A, sequence logo plots at right side of individual panels), which revealed that phosphorylation sites downregulated early and transiently by PI3K-TAK inhibition (cluster #6) are enriched in acidophilic residues, while the sites having the same trend upon PI3K-MEK inhibition do not show that enrichment. Overrepresentation analysis of known kinase motif sequences confirmed that cluster #6 is significantly enriched in sites located in acidophilic motifs: the casein kinase I substrate motif (FDR p-value: 3.03 e-5), the beta-adrenergic receptor kinase substrate motif (FDR p-value= 3.13 e-10) or the pyruvate dehydrogenase kinase substrate motif (FDR p-value= 0.004)

**Figure 3.**
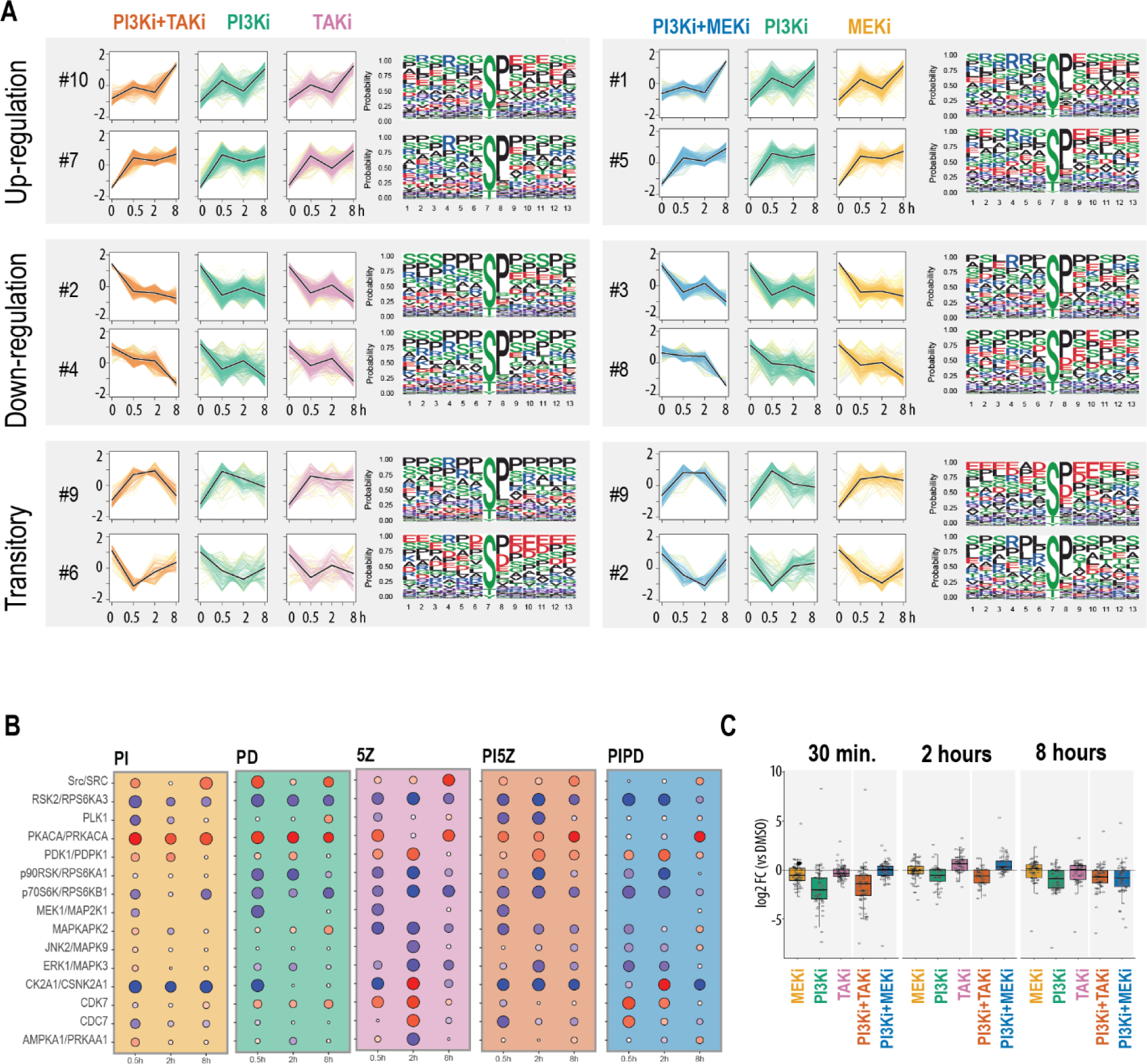
Phosphoproteomics temporal profiling of kinase inhibitor responses reveals downstream effects upon combination of treatments. (A) Temporal profiles after treatment with each kinase inhibitor or combination of them. Clusters were calculated using differentially regulated phosphosites for the synergy treatments (in at least one time point), separately. Only clusters with enriched kinetic trends are plotted. Following each trend in the synergies (k-means cluster), the temporal profile of the same phosphosite in the individual treatments is shown. Black lines indicate the centroid of the distribution. The phosphorylation intensity of each phosphosite used for clustering is the result of the average of biological replicates and then z-transformed across time points. On the right of each cluster profile, the amino acid logo sequence is shown, where the color indicates whether the amino acid is acidic (red), basic (blue), hydrophobic (black), neutral (purple), or polar (green). (B) Kinase activity inference using PTM-SEA. The color of the points indicates the fold-change (red: activation of the kinase, blue: inhibition of the kinase). Size of the point indicates the significance (-log10 q.value). (C) Boxplot showing regulation of CSNK2A1 known substrates (from PTM-SEA db) in each treatment and time point. Each point corresponds to the log2 fold change of a phosphorylation site in treatment versus its corresponding DMSO control.

To further explore these similarities and divergences, we evaluated kinase activity enrichment by PTM-SEA ^22^. There, we observed a consistent down-regulation of the PI3K/AKT/mTOR pathway in all treatments, indicated by the reduction of activity of RPS6KA1/3 (Figure 3B). However, in this analysis, the casein kinase 2 CSNK2A1 showed a dramatic differential regulation between individual and combined treatments. In individual treatments, only PI3Ki has a consistent inhibitory effect on CSNK2A1. However, this effect is only maintained in the combined treatment with TAKi, whilst the inhibitory effect is absent when combining PI3Ki with MEKi (Figure 3C). In summary, the kinome analysis shows that no new kinases are modulated in response to the drug combinations, with all estimated changes in kinase activity being observed also in at least one of the individual drugs.

### Drug combination response profile: Insights into overlaps and differences between drug combinations and single drugs

Having established that phosphorylation responses showed both overlaps and differences between conditions, we constructed the “response profile” of each synergy. The response profile refers to the quantification of all potential combinations of responses observed for two individual drugs and their combination. The term “response” denotes the qualitative changes in phosphorylation levels in the treated conditions relative to the DMSO control, measured by logFC values. From a theoretical perspective, with each phosphosite having the potential to increase, decrease, or remain unchanged across three conditions, a total of 33 = 27 distinct response patterns can be considered. These 27 response patterns were subsequently categorized into five primary classes (Figure 4A):

- **Combination-specific changes:** Differentially phosphorylated phosphosites (DPPs) that are only found in the combined inhibition or that have a response opposite to what is seen for each of the two single drugs.
- **Single drug-driven changes:** DPPs that change in the combination, but the response can be attributed to one of the single drugs.
- **Concerted changes:** DPPs that respond similarly across all conditions
- **Counterbalanced changes:** DPPs that show opposite responses between the two single drugs and are reflected as zero-sum changes in the combination.
- **Single drug-specific changes:** DPPs that are exclusively observed in one of the two individual drugs and not in any other condition.

**Figure 4.**
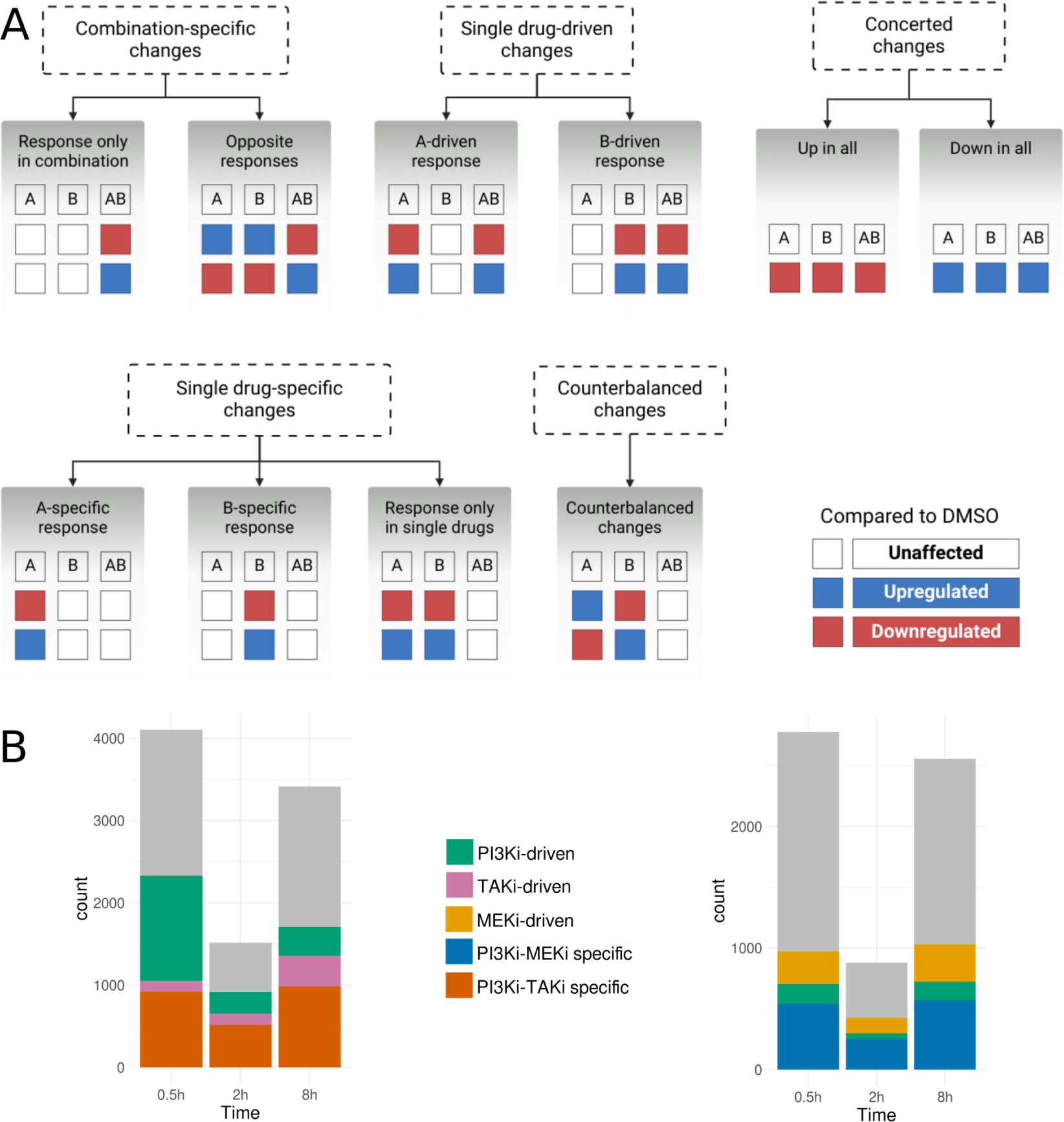
Response profiles categorize changes based on their driver drug (single versus combination). (A) Qualitative characterization of response classes in drug combinations and single drugs. Each molecular entity (i.e., gene or phosphosite) can have three states compared to the baseline: upregulated (blue-colored boxes with upward arrow), downregulated (red-colored boxes with downward arrow), or unaffected (white boxes). The set of states in the two drugs and their combination can be used to characterize each molecular entity and assign it to five main response classes. A representative example of each class is given. (B) Quantification of each change type for the PI3Ki-MEKi and PI3Ki-TAKi synergistic combinations.

The response profiles were defined independently for each time point, and the combination-specific changes were further assessed for consistency over time. Here, only those DPPs that were consistent across all time points were considered. DPPs that were categorized differently at different time points were not included in this analysis and were instead studied by a clustering analysis, which allows a more detailed description of more complex temporal behaviors.

### Combination-specific & single drug-driven effects combine to reduce cell growth

As depicted in Figure 4B, approximately 50% and 34% of the total DPPs for the PI3Ki-TAKi and PI3Ki-MEKi treatments, respectively, exhibited consistent changes across all conditions. Given the crosstalk between the targeted pathways, these coordinated changes could suggest at least a partially overlapping response mechanism between the single drugs that is also retained in their combination. Furthermore, as previously hinted by the correlation analysis shown in Figure 2D, many of the changes observed in the two combinations can be traced back to at least one of the two drugs. Specifically, in the PI3Ki-TAKi combination, the dominant influence of PI3Ki is evident at the 0.5-hour mark, primarily driving changes in synergy. Over time, this effect narrows down to a limited set of phosphosites, with most responses being combination-specific. Conversely, in the PI3Ki-MEKi combination, PI3Ki-driven changes are less prominent, with the majority of single drug-induced changes attributed to MEKi. Interestingly, the PI3Ki-MEKi combination exhibits a higher number of leveled-out responses across all time points compared to PI3Ki-TAKi. Specifically, at 0.5, 2, and 8 hours, PI3Ki-MEKi shows 122, 44, and 46 leveled-out changes, while PI3Ki-TAKi has only 22 leveled-out changes at 0.5 and 2 hours and 28 at 8 hours. Given the common use of PI3Ki in both combinations, these findings suggest that MEKi may counteract some of the effects of PI3Ki. When comparing the two combinations that synergistically inhibit growth, it appears that PI3Ki-TAKi has a higher number of combination-specific changes, with 2422 PI3Ki-TAKi-specific DPPs, compared to only 1368 combination-specific DPPs for PI3Ki-MEKi. A closer look at the overlap between the combination-specific changes of the two synergies showed that 338 of the combination-specific DPPs are shared between the two synergies. At the same time, the overlap between time points was limited to only a few common phosphosites, as most phosphosites were transiently regulated.

Next, we explored the combination-specific changes to identify whether these regulatory effects could represent candidate mechanisms that lead to the synergistically reduced cell growth observed for the two combinations. Both drug combinations exhibited combination-specific changes in several proteins related to cell survival. More specifically, in the PI3Ki-TAKi combination, RB1 exhibited a strong upregulation (logFC = 4.8, FDR p-val = 6.86E-10) at Ser-838, a site phosphorylated by p38, and induced the inhibitory interaction between RB1 and E2F1/2. Another interesting finding was the differential regulation of four phosphosites in the FOXO3 protein, two of which are known to affect its activity. Specifically, Ser-7 of FOXO3, also regulated by p38, leads to the nuclear accumulation of FOXO3. This particular phosphosite was uniquely upregulated by PI3Ki-TAKi treatment (logFC = 1.3, FDR p-val = 4.52E-05) at 0.5 hours. Moreover, FOXO3’s activating phosphosite Thr-32 (M2) was found with strong upregulation at 0.5 (logFC = 1.6, FDR p-val = 0.02) and 8 hours (logFC = 2.9, FDR p-val = 0.02). Interestingly, while Thr-32 upregulation at 0.5 hours can be attributed to PI3Ki, the upregulation of this site is retained and amplified only in PI3K-TAKi at 8 hours. Taken together, the joint inhibition of PI3K and TAK results in reduced cell growth ^12^, where phosphorylation changes from the PI3K and TAK1 pathways converged at the stress kinase regulation (p38) and cell fate regulation through the PI3Ki-driven regulation of FOXO3.

For PI3Ki-MEKi, fewer of the proteins, identified in our analyses with phosphorylation sites regulated in a combination-specific manner, have been reported to be involved in cell fate decisions, making it difficult to hypothesize mechanisms by which combination-specific phosphorylation changes could affect the cell’s physiological state. Potentially, the reduced cell growth observed with PI3Ki-MEKi could result from the cumulative effect of the two inhibitors on the same proteins, rather than from additional proteins that were differentially regulated when the two inhibitors were combined. Additionally, many of the PI3Ki-MEKi-specific sites were associated with controlling the protein’s intracellular localization or its molecular interactions but not directly with their activity. Nevertheless, some PI3Ki-MEKi-specific phosphorylation events are interesting to consider in relation to the cellular effects of the combination of these two inhibitors. The Ser-729 site of BRAF, which controls its enzymatic activation and is associated with cell growth inhibition, was upregulated uniquely in PI3Ki-MEKi at 8 hours. Ser-750, which inhibits BRAF-RAF1 interaction, was downregulated for up to 2 hours and could be attributed to the MEKi inhibitor that exhibited the same regulation. In addition, another activating phosphosite, Tyr-705, was upregulated at 8 hours. Additionally, two TFs were regulated in a PI3Ki-MEKi-specific manner. The cell fate regulator STAT3 appeared to be inhibited by the downregulation of the Ser-727 site, which activates the TF and inhibits apoptosis, at 0.5 and 2 hours. Ser-727 is regulated by several kinases *in vivo*, including JNK1, JNK2, and mTOR, whose activity changed in response to PI3Ki-MEKi treatment (Figure 3B). Lastly, FOXO3 displayed a similar regulation as in the other combination, with Thr-32 upregulation starting at 0.5 (logFC = 1.2, FDR p-val = 0.02) and retained at 8 hours (logFC = 3.4, FDR p-val = 6.62E-07).

In summary, the individual drugs and combinations negatively impacted several cell growth activators, aligning with their observed anti-growth properties ^12^. Notably, the observed synergistic growth inhibition in the combinations can be understood to arise from an aggregation of single drug-driven effects and specific changes in the phosphorylation of cell fate regulators. Furthermore, the effects of these combinations appear confined to their targeted pathways, with no indications of additional pathways being disrupted. This suggests that the synergistic growth inhibition can mostly be attributed to entities at the downstream convergence points of those pathways affected by the individual drugs rather than the dysregulation of new pathways. Taken together, these findings point towards growth arrest with a parallel apoptosis induction as evidenced by the activation of FOXO3 and BAD.

### Gene expression changes can be mostly attributed to combination-specific effects

To assess the temporal impact of kinase inhibition on gene transcription, we analyzed gene expression profiles at five time points: 1, 2, 4, 8, and 24 hours. Similar to the phosphoproteomics experiment, we employed DMSO and untreated samples as controls. The analysis was conducted using the limma-voom framework ^23^, with DMSO as the baseline at each time point, and declared a gene as differentially expressed (DEG) meeting an adjusted p-value threshold of 0.05 and an absolute logFC threshold of 1.

PCA analysis reveals that during the early time points (0 and 1 hour), all experimental conditions cluster together, an observation that was expected due to slower dynamics of transcription compared with phosphorylation events (Figure 5A). A notable divergence occurs in the 2-hour samples, particularly for MEKi and TAKi, which clustered closely until 8 hours, diverging only at the 24-hour time point. At the 24-hour time point, MEKi and PI3Ki-MEKi samples exhibited distinct clustering, separated from the rest of the samples (Figure 5A). Most of the variation captured by the first principle component appears to be related to time, while the second and third components capture mainly the influence of treatment. For a broader perspective, we investigated the correlation between the effects of single drugs and their respective combinations and found a strong positive Pearson correlation, with all correlation coefficients > 0.88 (Supplementary Figure 4), thereby suggesting that the effects of individual inhibitors are largely retained in the combination treatments.

**Figure 5.**
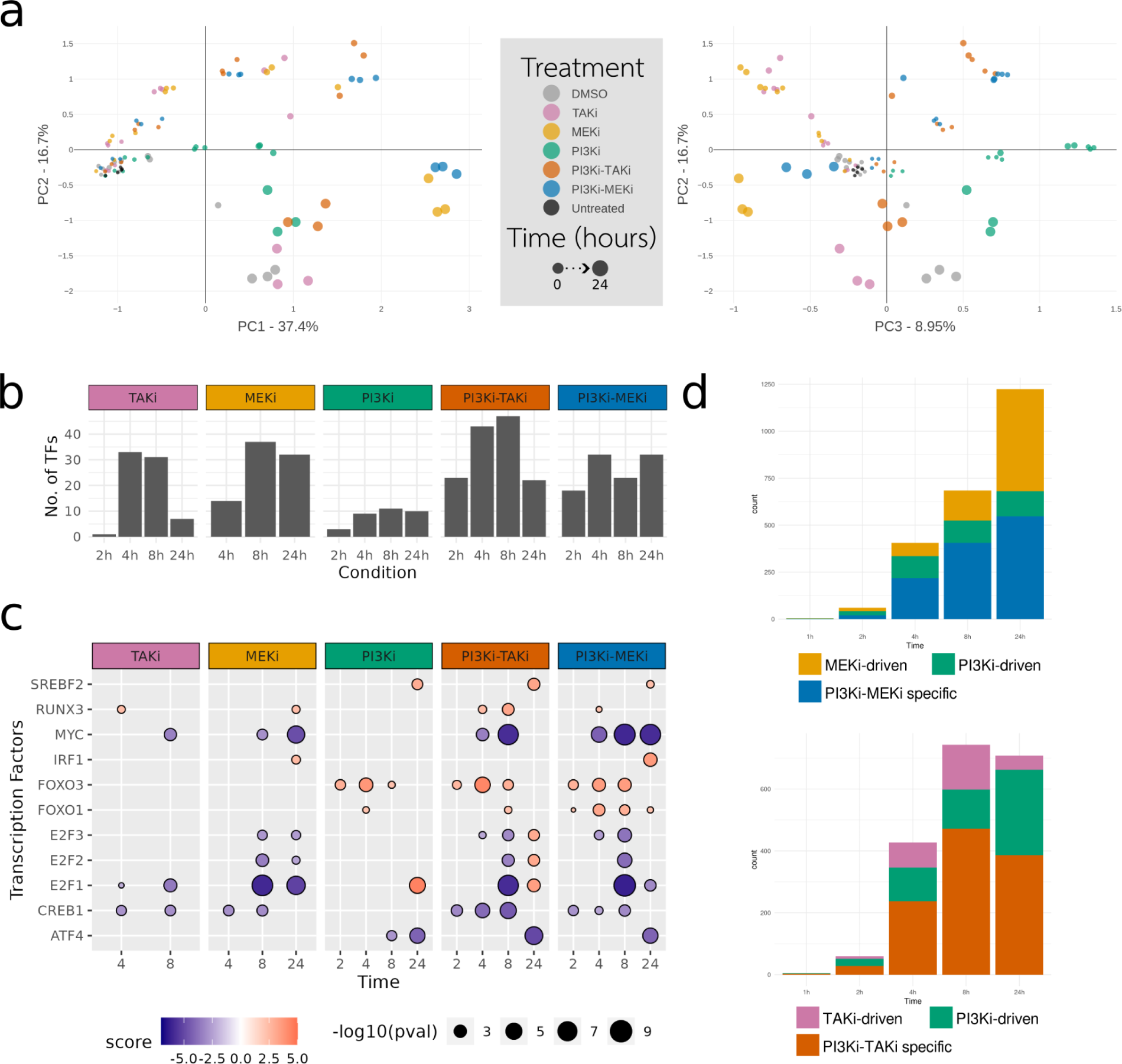
Transcriptomics analysis shows a high number of combination-specific changes and an altered activity of key cell fate-regulating transcription factors. (A) Principal Component Analysis of transcriptome profiles of samples treated either with DMSO or with each drug combination, and time zero (non-treated samples). (B) Number of transcription factors with an estimated differential activity per drug and time point. Activities were estimated using a univariate linear model and the CollecTRI regulon. P-adjusted threshold 0.05. (C) Activities of transcription factors that displayed the highest and lowest estimated activities. (D) Quantification of each change type for the PI3Ki-MEKi and PI3Ki-TAKi synergistic combinations.

To identify regulators of differentially expressed genes (DEGs) in each condition, we conducted a transcription factor (TF) activity analysis, focusing on TFs with measured gene expression or protein abundance in our datasets for relevance (Figure 5B and Supplementary Figure 5). Changes in TF activities unique to combinations showed smaller magnitude and significance compared to changes shared with at least one single drug (Figure 5C and Supplementary Figure 5). Additionally, consistent with phosphoproteomics observations, our transcriptome findings indicate growth inhibition across all conditions, with reduced activity for TFs like CREB1 and MYC (Figure 5C), known for supporting proliferation and survival. Notably, both MYC and CREB1 showed more pronounced activity reduction in combinations compared to single drugs. The influence of PI3Ki on the PI3K/AKT/FOXO3 pathway is reflected in gene expression, with FOXO3 showing increased activity in PI3Ki and both combinations. Furthermore, the antagonistic interaction between PI3Ki and MEKi was evident in TF activities estimated from gene expression, as, for example, seen in the regulation of the E2F TF family (Figure 5C). Lastly, the PI3Ki-MEKi-specific inhibition of STAT3 activity at early time points (Supplementary Figure 5) was mirrored by our findings concerning its phosphorylation status, where its activating phosphorylation site was downregulated at 0.5 and 2 hours. However, no change in its activity or phosphorylation was observed at later time points.

For transcriptomics, we performed separate clustering for each drug by an unsupervised k-means approach. For each cluster, we conducted a Gene Ontology (GO) term overrepresentation analysis. Notably, GO terms linked to cell cycle progression and phase transitions were significantly overrepresented in late downregulated clusters (peaking at 8 and/or 24 hours) across all drugs. Interestingly, the median temporal trends of most cell cycle-related clusters aligned with those of the DMSO control at 24 hours. This suggests that cell cultures, even in the absence of kinase inhibitors, may exhibit reduced cell cycle activity, potentially influenced by DMSO or reduced proliferation due to contact inhibition. GO terms related to RNA processing (GO:0006396), ribosome biogenesis (GO:0042254), and rRNA processing (GO:0016072) were overrepresented in temporal clusters that displayed an early onset of downregulation up to 8 hours for PI3K-TAKi and 24 hours for PI3K-MEK. The same terms were the only GO terms overrepresented among the combination-specific DEGs at 8 hours (Supplementary Tables 1 and 2), with all genes annotated to these GO terms being downregulated. Among genes annotated to ribosome biogenesis, we observed the downregulation of several subunits of RNA Pol II and III. The expression of these genes is known to be influenced by mTOR and MYC ^24^, which aligns with the reduced activity estimated for these two factors. Furthermore, genes annotated to macroautophagy and terms related to its positive regulation were overrepresented in early onset clusters of PI3Ki and the two combinations (Supplementary Figure 7). Although the phosphorylation and expression levels of autophagy regulators and effectors point to a PI3K-driven induction of autophagy from early time points up to 8 hours, the expression landscape at 24 hours suggests that autophagy is not maintained at later time points (Supplementary Figure 7). Alternatively, the time point when the autophagy manifests may both be later and last longer than the time points when transcripts encoding protein effectors of the cellular response are observed with changed levels.

The intrinsic apoptotic signaling pathway term (GO:0097193) was overrepresented only in the two combinations at 24 hours. The only single drug that had apoptosis-related terms overrepresented was TAKi, where the positive regulation of extrinsic apoptotic signaling pathway (GO:1902043) was overrepresented at 2 hours and the negative regulation of the same pathway (GO:2001237) was overrepresented at 24 hours. In line with our phosphoproteomic data indicating the induction of apoptosis via the activation of FOXO3, we observed that the apoptosis and anoikis regulator BCL2L11 (Bim) was uniquely upregulated in both combinations. In PI3Ki-TAKi, its expression was upregulated at 4h. In PI3Ki-MEKi, BCL2L11 displayed a more consistent upregulation at both earlier and later time points (2, 4, and 24 hours). FOXO3 is known to cooperate with RUNX3 to induce BCL2L11 expression in gastric adenocarcinoma ^25^. Such FOXO3-RUNX3 cooperativity could potentially play a role in increased BCL2L11 expression in both combinations, with both TFs having a predicted increased activity (Figure 5C). The activation of FOXO family members plays a diverse role in gastric cancer, acting through the induction of both apoptosis and cell cycle arrest ^26^. More specifically, FOXO4, uniquely activated by the two combinations, has been reported to lead to the induction of cell cycle arrest ^26^. Additionally, the anti-apoptotic BCL2L12 was downregulated at 24h solely in PI3Ki-MEKi. Altogether, gene expression suggests the parallel induction of cell cycle arrest and apoptosis as also indicated by the phosphoproteomics data, in both drug combinations. The reduced growth appears to be mediated through the reduced activity of proliferation-promoting TFs (MYC, CREB1), increased activity of apoptosis-promoting TFs (FOXO3, FOXO4, and RUNX3), and the altered activity and/or expression of cell cycle arrest regulators (CDKN1A, BCL2L11, CEBPA, SIRT4).

### Effect of drug combination on metabolism: Nucleotide metabolism and tRNA biosynthesis are downregulated only in the combinations

Among the combination-specific DEGs of both combinations, several were associated with key components of gastric cancer-promoting metabolic processes. Using gene-metabolic pathway associations from the genome-wide metabolic model (Human-GEM v 1.12.0) ^27^, the combination-specific DEGs could be mapped to two main metabolic processes: nucleotide metabolism and aminoacyl-tRNA biosynthesis. GO terms related to the same processes were also enriched in late downregulated clusters in the two combinations (Figure 6A).

**Figure 6.**
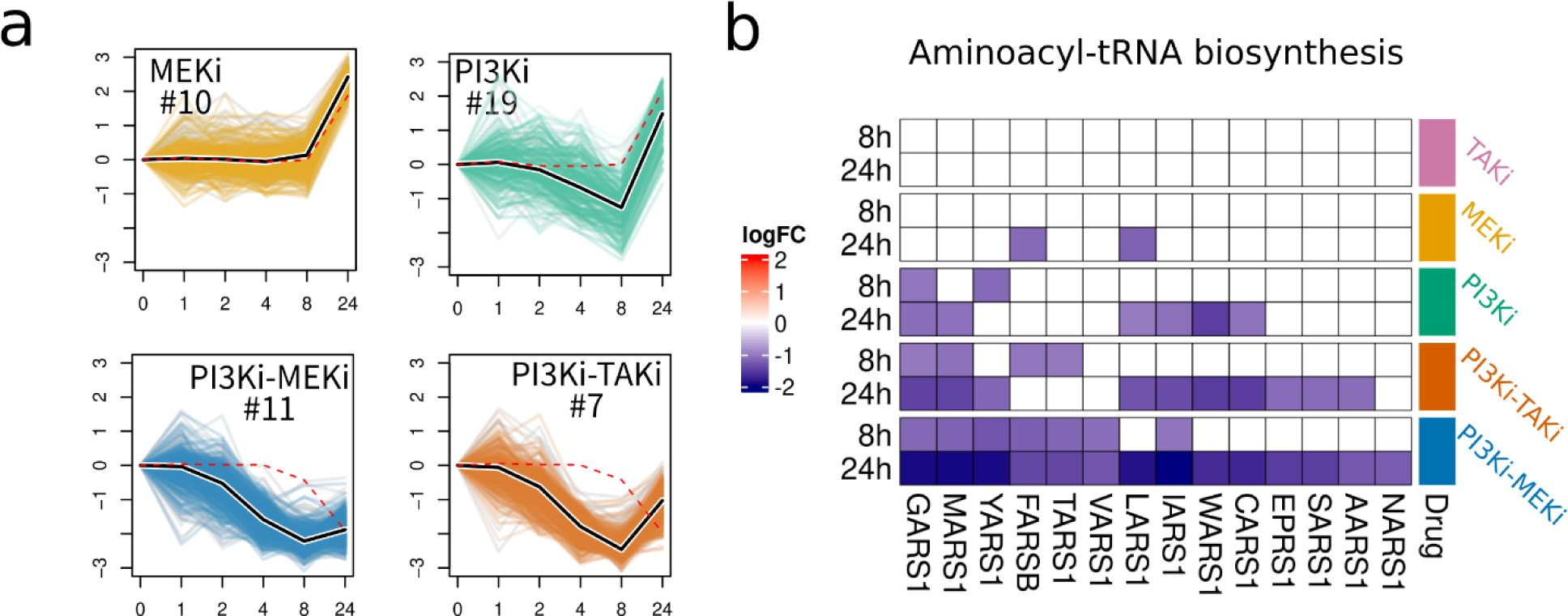
Nucleotide metabolism and aminoacyl-tRNA biosynthesis are downregulated in response to the drug combinations. (A) Gene expression temporal clusters with a significant enrichment (FDR p-value < 0.01) of nucleotide metabolism-related Gene Ontology terms. Black lines indicate the centroid of the distribution and red dash lines indicate the centroid of the distribution for the same genes in untreated conditions (DMSO). Gene expression levels used for clustering is the result of the average of the biological replicates and then z-transformed across time points. (B) Heatmap of the logarithmic fold change (logFC) of the differentially expressed aminoacyl synthase enzymes at 8 and 24 hours. Earlier time points are not shown as none of the enzymes were differentially expressed.

The downregulation of nucleotide metabolism by the PI3Ki-TAKi combination was also supported by phosphorylation changes and these were not observed with the single inhibitors. CAD, a ‘fusion’ gene that encodes key enzymes involved in the pyrimidine metabolic pathway, displayed downregulation of its activating phosphosite Ser-1859 at 2 and 8 hours uniquely in PI3Ki-TAKi. Ser-1859 is a target of p70S6K; therefore, its downregulation could be a downstream effect of mTOR inhibition. Additionally, TAKi-driven CAD Ser-1406 phosphorylation, which is known to block the phosphorylation of another activating site of CAD, was consistently upregulated. PDHA1, involved in pyruvate metabolism, exhibited upregulation of inhibiting sites Ser-232 and Tyr-289 in PI3Ki-TAKi. Similarly, genes associated with nucleotide metabolism were mostly downregulated in the two combinations, a behavior that could not be attributed to any of the single drugs (Figure 6A and Supplementary Figure 6). For tRNA biosynthesis, we observed a significant downregulation of most aminoacyl tRNA synthetase enzymes (Figure 6B). In the context of senescence, mTOR has been found to regulate tRNA biogenesis and specific aminoacyl-tRNA synthetases, namely LARS and YARS ^28^, both of which were among the most strongly downregulated aminoacyl-tRNA synthetases. In a colorectal cancer cell line, the downregulation of LARS is reported to be associated with the downregulation of E2F1-modulated proliferation genes and cell cycle arrest ^28^, both of which are observed in our results as well. An additional potential mechanism for the downregulation of aminoacyl tRNA synthetases could be attributed to the stress-induced ATF4 transcription factor, which activates the expression of the majority (16 out of 20) of these enzymes ^29^. Multiple mechanisms, including ubiquitination, phosphorylation, and transcription, regulate the activity of ATF4. When assessing the phosphorylation status of ATF4, it appears that the protein is stabilized via the downregulation of its Ser-248 site, which contributes to protein destabilization and degradation. For the combinations, Ser-248 is strongly downregulated at 2 hours (*PI3Ki-TAKi:* logFC = -3.8 and FDR p-val = 8.7*10^-9^ and *PI3Ki-MEKi:* logFC = -3.5 and FDR p-val = 4.6*10^-9^) and 8 hours (*PI3Ki-TAKi:* logFC = -3.5 and FDR p-val = 8.1*10^-7^ and *PI3Ki-MEKi:* logFC = -3.4 and FDR p-val = 1.2*10^-5^). For the single treatments, Ser-248 is upregulated at 0.5 hours post-treatment with MEKi (logFC= 1.6 and FDR p-val = 4.6*10^-5^) and TAKi (logFC= 1.5 and FDR p-val = 0.004), and downregulated at PI3Ki but only at 2 hours (logFC= 2.8 and FDR p-val = 7.5*10^-6^). At the transcriptome level, ATF4 expression levels are downregulated in all conditions at 8 hours, but this downregulation is retained only by the two drug combinations at 24 hours when also its activity is predicted to be reduced. Inhibiting ATF4 activity has been an attractive treatment option as high ATF4 activity has been associated with poor prognosis in gastric cancer ^30^ and its increased expression provides tumors the ability to adapt to microenvironment-related stress ^31^. Among the strategies for its downregulation is the targeting of upstream eukaryotic translation initiation factors (eIFs) kinases, which reduces its translation ^31^. In our dataset, we observed a combination-specific downregulation of the expression levels of multiple eukaryotic translation initiation factors (eIFs), which play a critical role in regulating the initiation stage of protein synthesis. Notably, the activity of eIFs is governed by both the PI3K/AKT and MAPK pathways ^30^. This could explain why only the drug combinations inhibiting both these pathways downregulate eIF, while such downregulation was not observed with the single drugs. Downstream propagation of eIFs is likely to result in downregulation of ATF4, thereby impairing amino acid biosynthesis at 24 hours post-treatment, both events having been reported to subsequently result in increased oxidative stress, cell cycle arrest, induction of apoptosis, and delayed tumor growth ^31^.

## Discussion

We have employed a multi-omics approach combining phosphoproteomics and transcriptomics to evaluate the temporal effects up to 24 hours of combining kinase inhibitor treatments of a gastric adenocarcinoma cell line. Previously, *in silico* model-based simulations ^12^ revealed the potential synergy between PI3K and MEK or TAK inhibition for this type of cancer with subsequent experimental validation of these drug synergies ^12^. However, models, such as the one used in our previous work ^12^, do not provide precise insights into molecular processes underlying the regulatory mechanism of synergy or how this may result in an improved therapeutic effect. Phosphoproteomics offers insights into the first line of action of kinase inhibitors since their effects, are exerted at the protein level. Protein phosphorylation events are central to the modulation of signaling pathway kinase cascades and are commonly very fast. Our data validate the expected targets of each treatment but also inform about the connectivity between them. While previous studies have examined the synergistic effect of targeting PI3K and MAPK pathways ^33–38^, these studies are mostly limited to the targeted measurement of (phospho)proteins or gene expression, without exploring the mechanism of action of the drug combination in a systemic way or in multiple biological levels. This integrated, multi-omics analysis allowed us to track the responses upon drug combination treatment, from the fast inactivation of the targeted kinases to the downstream effect on the targeted pathways and the subsequent effect on the transcriptional changes (summarized in Figure 7).

**Figure 7.**
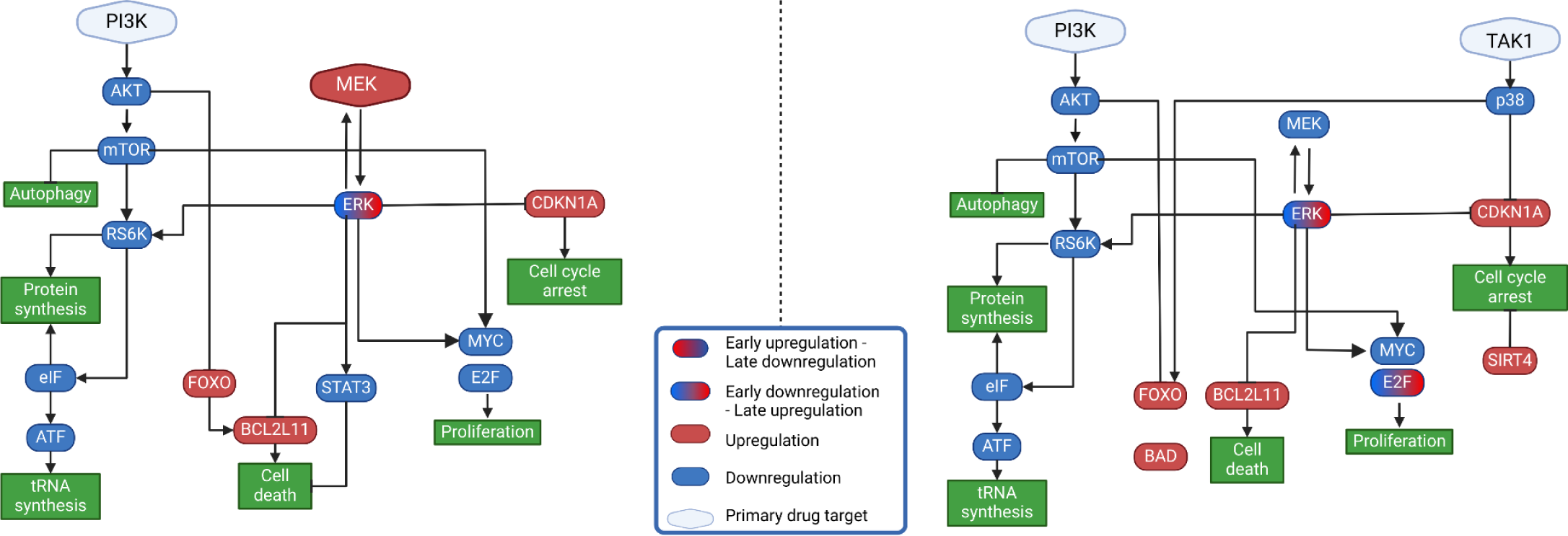
Proposed mechanisms of action of drug combinations up to 24 hours post-treatment. (A) PI3Ki-TAKi mechanism, and (B) PI3Ki-MEKi mechanism The proposed models show changes in phosphorylation and/or gene expression in response to the drug combinations, including combination-specific changes and those driven by the individual drugs. Green rectangles represent biological processes in which the dysregulated entities are involved.

Overall, the various analyses indicate that the synergistic effect of the combinations is driven by changes at the points of convergence of the two pathways, but not at the kinase level. This is particularly evident from the phosphoproteomics where there are no changes in the regulation or the activity of kinases that cannot be attributed to either or both single drugs. The pathways targeted by the chosen inhibitors have a considerable degree of cross-talk and constitute some of the main tumorigenic pathways that regulate critical cancer-promoting processes and cell fate decisions ^36^. Consequently, even in the absence of modulation of additional kinases, when the two drugs are combined, we observe changes in entities regulated by both pathways at downstream regulatory levels, namely phosphosites, transcription factors, and genes. This suggests that the synergistic growth reduction is a consequence of the blockade of alternative ways that would otherwise be employed to counteract the effects of the single drugs. An example of such effects is the combination-specific regulation of the apoptosis-promoting BCL2L11, where FOXO3 and RUNX3 are known to cooperatively induce BCL2L11 expression in gastric adenocarcinoma ^25^. In our dataset, we observe the PI3Ki-driven activation of the FOXO TF family and the MEKi and TAKi-driven RUNX3 activation that combine to induce the gene expression of BCL2L11. This can also be seen in combination-specific DEGs that are intricately linked to pivotal components of processes known to fuel gastric cancer progression. Notably, nucleotide metabolism and aminoacyl-tRNA biosynthesis pathways stood out as particularly affected with many of the main genes involved in the two processes being downregulated. Aminoacyl-tRNA biosynthesis is directly involved in the progression of gastric cancer, and its targeting has been proposed as a viable treatment strategy ^29,39^. Similarly, sustained and aberrant nucleotide metabolism is a critical component in cancer development as it constitutes a cancer dependency independently of cancer type, tissue of origin, or driving molecular alteration ^40^. The regulation of those processes is under the influence of both PI3K/AKT and MAPK pathways. As a result, we observe responses unique to the two combinations that start from the downregulation of eIF factors, to the reduced activity of ATF4 and the halted aminoacid biosynthesis at 24 hours after combination treatment. Combination therapies have been proposed as an option for targeting nucleotide metabolism in cancer ^40,41^, and the combinatorial targeting of PI3K-MAPK pathways presented in this study could be further explored as a potential treatment against those processes.

In addition to the synergistic changes, we also shed light on the additive or antagonistic interactions between the single drugs. As mentioned above, we observe a high overlap between the single drugs and their combinations, through our response profile analysis. We observed that almost half of the phosphorylation changes at any given time were the same, both for the combinations and their single drugs, which could suggest a shared basic layer of drug response mechanism for all drugs and their combinations. It was also previously shown that drugs modulating cell fate decisions and inducing cell death do so by a similar mechanism resulting in a distinct cell death signature that could inflate the levels of similarity between the mechanism of actions of different drugs ^37^. However, since this work compares single drugs and their combinations, it is assumed that the cell death signature only partially explains the high correlation observed. Interestingly, we also found several instances where MEK inhibition can counteract the effect of PI3K inhibitors both in phosphorylation status and gene expression. CSNK2A1 activity is one instance where we observed such an antagonistic interaction, as our data reveals that upon PI-103 treatment on its own, or in combination with 5-z-7-oxozeaenol, CK2A is rapidly inhibited at 30 minutes of treatment. However, when PI-103 is combined with the MEK inhibitor, such inhibition is abrogated. CSNK2A1 inhibition has been previously identified as a complementary target to enhance the effectiveness of PI3K inhibition in colorectal cancer ^39^. Interestingly, it has been also described that CSNK2A1 blockade can potentiate the antiproliferative effects of BRAF and MEK inhibition in BRAF-mutated cancers ^40^. Altogether, our findings are in line with the work of Olsen et al. on glioblastoma cells that connects CSNK2A1 activity with the mTOR/AKT/PI3K axis, and a negative feedback between CSNK2A1 with MEK/ERK regulation ^41^. This would suggest that MEKi could counteract the inhibitory effect of PI3Ki on CSNK2A1. On the other hand, this suggests that it should be interesting to investigate a potential synergistic effect of inhibiting CSNK2A1 in combination with MEK inhibition.

Lastly, it was interesting to observe that for both of the combinations, their molecular response hallmarks are indicative of cytostatic as well as cytotoxic effects at the cellular level. While many studies often concentrate on differentiating between cytotoxic and cytostatic drugs, there is no well-defined molecular distinction between the two, with many of the drugs exhibiting properties of both, depending on the context ^42^. In both experimental and clinical scenarios, cytotoxic substances can also induce cell stasis when administered in low doses or when cells are resistant to apoptosis. Conversely, cytostatic compounds can trigger apoptosis in cancer cells that are in states other than quiescence ^42^. Additionally, prolonged cell stasis in response to cytostatic compounds can lead to cell death ^42^. In our study, however, it seems unlikely that we could observe this effect due to the short time frame of our investigation. In our experimental design, we used unsynchronized cell populations as this scenario resembles real-life conditions. This decision could explain why we find evidence that supports both a stasis of cell growth and induction of cell death in response to both of the combinations, as cells in different cell cycle states could be responding differently. In PI3K-MAPK co-targeting in thyroid cancer cell lines others have observed that the cells exhibited cell cycle arrest that led to apoptotic cell death but only after 24 hours ^38^. However, our main aim remains to describe the molecular events that lead to the synergistic reduction of cell growth and other types of experiments should be employed to characterize a compound or a combination as either or as both. Additionally, it was previously shown that responses in PI3K-MAPK double inhibition were dependent on the mutational status of the tested cell lines ^43,44^, and therefore follow-up experiments might be required to explore the effect of mutations to combination responses either in a panel of cell lines or in more complex experimental models, such as organoids.

In conclusion, our study provides a comprehensive view of the molecular events occurring in AGS cells when exposed to PI3K, TAK1, and MEK inhibitors both individually and in combination. The observed changes in phosphorylation, transcription factor activity, and gene expression collectively support altered cell fate decisions and metabolic dysregulation as central mechanisms underlying the synergistic effects of these combinations.

## Materials and Methods

### Experimental setup

AGS (human gastric adenocarcinoma, ATCC, Rockville, MD) were grown in Ham’s F12 medium (Invitrogen, Carlsbad, CA) supplemented with 5% fetal calf serum (FCS; Euroclone, Devon, UK), and 10 U/ml penicillin-streptomycin (Invitrogen). Chemical inhibitors PI-103, 5Z: (5Z)-7-oxozeaenol, and PD: PD0325901 (all Merck) were solved in DMSO at stock concentrations of 20 mM.

### Raw data production & analysis

#### Transcriptomics

##### Sample preparation

Cells (0.5 *10^6^) were seeded in 6-well plates, reaching 80-90 % confluency after 24 hrs. Then inhibitors PI (0.70 µM), 5Z (0.50 µM), and PD (0.035 µM) were added (single or in combination), and cells were further incubated for 1, 2, 4, 8, and 24 hrs. One 6-well plate was prepared for each time point including 0 hour untreated control plate. Subsequently, media was removed, and cells were washed once with PBS (37 °C) and 350 µl RNA lysis buffer (Qiagen, Buffer RLT

Plus, Cat. No. 1053393) added. Cell lysates were further transferred to liquid N2/ -80 °C freezer before RNA purification according to Qiagen protocol (www.Qiagen.com; AllPrep DNA/RNA/miRNA Universal Kit, Cat. No. / ID:80224).

##### Library construction and sequencing

RNA concentration was measured using Qubit® RNA HS Assay Kit on a Qubit® 3.0 Fluorometer (Thermo Fisher Scientific Inc., Waltham, MA, USA). Integrity was assessed using Agilent RNA 6000 Pico Kit on a 2100 Bioanalyzer instrument (Agilent Technologies, Santa Clara, CA, USA). RNA sequencing libraries were prepared using the Illumina Stranded mRNA prep ligation kit (Illumina, San Diego, CA, USA) according to the manufacturer’s instructions. The final libraries were purified using the AMPure XP (Beckman Coulter, Inc., Indianapolis, IN, USA), quantitated by qPCR using KAPA Library Quantification Kit (Kapa Biosystems, Inc., Wilmington, MA, USA), and validated using Agilent High Sensitivity DNA Kit on a Bioanalyzer (Agilent Technologies, Santa Clara, CA, USA). The size range of the DNA fragments was measured to be in the range of app. 200-1000 bp and peaked at around 300 bp. Before sequencing, the libraries were quantified (KAPA Library Quantification Kit (Illumina/ABI Prism), normalized, and pooled. Quantitated libraries were further diluted to 2.5 nM and subject to clustering by a cBot Cluster Generation System on four HiSeq4000 flow cells (Illumina Inc. San Diego, CA, USA), according to the manufacturer’s instructions. Finally, single-end read sequencing was performed for 64 cycles on an Illumina HiSeq4000 instrument, following the manufacturer’s instructions (Illumina, Inc., San Diego, CA, USA). FASTQ files were created with bcl2fastq V2.20 (Illumina, Inc., San Diego, CA, USA).

##### Raw data processing

The quality of the produced FASTQ files was controlled with fastqc (version 0.11.9) and then filtered and trimmed by fastp (version 0.20.0). Trimmed sequences were aligned to the genome reference using STAR (version 2.7.3) and quality metrics were extracted with picard CollectRNASeqMetrics (version 2.21.5). Transcript counts were generated using quasi alignment with Salmon (version 1.3.0) to the GRCh38 transcriptome reference sequences. Transcript counts were imported into the R statistical software and aggregated to gene counts using the tximport (v1.14.0) Bioconductor package for downstream statistical analysis.

#### Phosphoproteomics

##### Sample preparation

Cells (4 *10^6^) were seeded in TC175 flasks in a total volume of 10 ml medium with 5% FCS. One TC75 flask was prepared for each time point, plus one additional flask for DMSO for 8 hrs. After leaving cells in the incubator overnight for 20 hrs, inhibitors PI-103 (0.70 µM), 5-z-7-oxozeaenol (0.50 µM), and PD0325901 (0.035 µM) (single or combinations) were added to each TC75-flask except flasks to be harvested at 0 hrs. Cells were then incubated for 0.5, 2, and 8 hours. Subsequently, media was removed, cells washed once with PBS and trypsinized using 3 ml trypsin/EDTA, left until detachment, typically around 3-5 minutes. 7 ml media with 10% FCS was added, cells collected, pellet (1500 RPM for 5 minutes, 4 °C) and washed with 5 ml PBS (4 °C) supplemented with phosphatase inhibitors (1mM NaF, 1mM beta-glycerol phosphate and 5mM Sodium Orthovanadate) and additional pellet (1500 RPM for 5 minutes, 4 °C). The supernatant was removed and the pellet resuspended in 1 ml cold PBS. Cells were transferred to a 1.5ml Eppendorf tube, pellet (1500 RPM, 5 min, 4 °C), PBS removed and the cells were immediately frozen in liquid nitrogen before being transferred to a -80 °C freezer.

##### Sample preparation for MS analysis

Snap-frozen cell pellets were lysed in 300 µl of boiling lysis buffer containing 5% SDS, 100 mM TrisHCl pH 8.5, 5 mM TCEP, and 10 mM CAA for 10 minutes at 95ᵒC. Samples were further homogenized by sonication with a probe for 45 seconds (1 sec ON, 1 sec OFF, 30% amplitude). Protein concentration was measured by BCA. Afterward, samples were digested overnight using the PAC protocol ^45^ implemented for the KingFisher robot as described previously ^46^. Samples were acidified after digestion to a final concentration of 1% trifluoroacetic acid (TFA) and peptides were loaded onto Sep-Pak cartridges (C18 1 cc Vac Cartridge, 50 mg - Waters). Eluted peptides from the Sep-Pak were concentrated in a Speed-Vac, and 150 µg of peptides (measured by A280 Nanodrop) were used for phospho-enrichment. Phosphoenrichment was performed as described previously ^46^ using 20 µl of TiIMAC-HP beads (MagResyn). Eluted phosphopeptides were acidified with 10% TFA to pH <3, filtered through MultiScreenHTS HV Filter Plate (0.45 µm, clear, non-sterile) for 1 minute at 500 g and loaded into Evotips for further MS analysis.

##### LC-MSMS analysis

Samples were analyzed on the Evosep One system using an in-house packed 15 cm, 150 μm i.d. capillary column with 1.9 μm Reprosil-Pur C18 beads (Dr. Maisch, Ammerbuch, Germany) using the pre-programmed gradients for 60 samples per day (SPD) for phospho-proteome samples. The column temperature was maintained at 60°C using an integrated column oven (PRSO-V1, Sonation, Biberach, Germany) and interfaced online with the Orbitrap Exploris 480 MS. Spray voltage was set to 2.0 kV, funnel RF level at 40, and heated capillary temperature at 275°C. Full MS resolutions were set to 120,000 at m/z 200 and full MS AGC target was 300% with an IT of 45 ms. Mass range was set to 350−1400. A full MS scan was followed by a DIA scan comprising 49 windows of 13.7 Da with an overlap of 1 Da, scanning from 472 to 1143 Da for phospho-proteome and 361 to 1033 Da for total proteome. Resolution was set to 15,000 and IT to 22 ms. The normalized collision energy was set at 27%. AGC target value for fragment spectra was set at 1000%. All data were acquired in profile mode using positive polarity and peptide match was set to off, and isotope exclusion was on.

##### Raw data processing

Raw files were searched in Spectronaut (v16) using a library-free approach (directDIA). Carbamidomethylation of cysteines was set as a fixed modification, whereas oxidation of methionine, acetylation of protein N-termini, and, in phospho-proteomics samples phosphorylation of serine, threonine, and tyrosine, were set as variable modifications. Homo sapiens FASTA database (UniProtKB/Swiss-prot 21,088 entries) and a common contaminants database were used for directDIA search. For phospho-proteomics samples, the PTM localization cutoff was set to 0.75. Cross-run normalization was off and Data Filtering was set to Qvalue. Phospho-peptide quantification data was exported and collapsed to site information using the Perseus plugin described in Bekker-Jensen et al ^47^. Phosphosite data sets were processed using R (v3.6.2). Data was log2 transformed and three valid values in at least one experimental group were required to preserve the phosphosite for further analysis. Data was normalized to remove experimental bias due to sample handling using the Loess method. Imputation of missing values was performed using the data analysis pipeline of Prostar (v 1.18.4) ^48^. Imputation of missing values was performed in two steps: first partially observed values (i.e., values missing within a condition in which there are valid quantitative values) were imputed using the function slsa from the DAPAR package, and second, values missing in an entire condition were imputed using the detQuant function (quantile=1, factor=1) also from DAPAR package.

### Computational and statistical analysis

Unless stated otherwise, the analyses were performed in RStudio (version 2022.02.2) with R version 4.2.3 (2023-03-15). Principal component analysis (PCA) was performed using the PCAtools package (version 2.8.0) ^49^ and visualized using Plotly (version 4.10.1) ^50^. Heatmaps were created using the ComplexHeatmap R package (version 2.12.0)^51^. The rest of the visualizations were created using ggplot2 (version 3.4.1)^52^ unless stated otherwise.

#### Differential expression analysis

For phosphoproteomics, differential site regulation was calculated when at least 75% of the values were valid (not imputed) in one condition of each pairwise comparison. Significantly regulated sites were calculated using a moderated two-sided t-test (in limma R package) setting the method of the lmFit function to “robust” and using Benjamini-Hochberg FDR correction. For transcriptomics, a differential expression analysis was performed using the limma R package (version 3.52.0) ^23,53^. Gene counts were transformed using the limma-voom workflow. Genes with zero or very low (i.e. CPM < 3) counts were omitted from the analysis. For each comparison, a linear model was fitted for each gene with each condition (i.e. treatment and time) as a variable. DMSO was used as a baseline for all contrasts. Differentially expressed genes were defined using a logarithmic fold change (logFC) threshold of 1 and an adjusted p-value threshold of 0.05.

#### Response profiles

To gain a deeper understanding of how the two single drugs contribute to the changes (i.e. Differentially phosphorylated phosphosites and Differentially expressed genes) found in the synergies, we developed a decision tree that classifies each DPP and DEG into one of predefined categories. This categorization was qualitative and was based on whether a phosphosite or gene was upregulated, downregulated, or unaffected compared to DMSO, without taking into account the intensity of changes. As each phosphosite/gene can have three types of changes (i.e. upregulated, downregulated, unaffected) in three conditions (Drug A, Drug B, and Combo A+B), a total of 27 possible categories can arise. These categories were further grouped to main categories denoting whether a change was:

- specific to one condition (i.e. combination-specific, DrugA-specific, DrugB-specific changes),
- driven by one of the two single drugs (i.e DrugA-driven, DrugB-driven changes),
- same in all conditions (i.e. Concerted changes)
- antagonistic between the two drugs (i.e. Counterbalanced changes).

To validate the ability of the decision tree to correctly classify each phosphosite and gene, a toy dataset with all possible change types was created.

#### Phosphoproteomics Temporal Clustering

Temporal profiles were obtained for each synergy (PI3K+TAK inhibition and PI3K+MEK inhibition). Only phosphorylation sites that were significantly regulated (q-value <0.05) in one time point were used for further analysis. Temporal profiles were obtained by z-scoring the log2 fold change at each time point of treatment versus DMSO and setting the time zero value as 0. Clustering was performed using the cluster_analysis function from multiclust package (v1.24.0) in R, setting a fixed number of 10 clusters, with k-means as the clustering algorithm and Pearson as distance measure.

#### Kinase activity signatures enrichment analysis

Kinase activity inference was calculated using single sample Gene Set Enrichment Analysis approach with PTMSigDB, also known as PTMSignature Enrichment Analysis (PTM-SEA)) ^22^. Log2 fold change at each time point for each treatment versus DMSO was used as input for PTM-SEA analysis. A minimum of five phosphosite per signature is required to calculate the enrichment score. Kinase signatures with a q-value < 0.01 were considered significant..

#### Phosphoproteomics motif analysis

Motif logos for each phosphoproteomics temporal cluster were plotted using ggseqlogo (v0.1). Motif overrepresentation analysis was performed in Perseus (v1.6.5.0) using a two-sided overrepresentation test (Fisher exact t-test) and using the overall measured phosphoproteome as a background.

#### Transcription factor activity inference & Overrepresentation analysis

The metasource CollecTRI was used as a Prior Knowledge Network (PKN) for the estimation of transcription factor activity ^54^. Bioactivity estimations were performed using the decoupleR package (version 2.2.0)^55^ and the univariate linear model (ulm) method was used. Only transcription factors with minimum 5 annotated target genes in the PKN were included in the analysis. Overrepresentation analysis for Gene Ontology terms was performed with the clusterProfiler package (version 4.4.1) ^56^. The total of measured genes in our transcriptomics dataset was used as a background for the analysis, and with an adjusted p-value threshold of 0.05.

### Data and code availability

The mass spectrometry proteomics data have been deposited to the ProteomeXchange Consortium via the PRIDE partner repository ^57^ with the dataset identifier XX. Raw transcriptomics data are available at XX. The code for all the presented analyses is available in https://github.com/Eirinits/AGS_synergies.git.

## Acknowledgements

The RNA library prep, sequencing and bioinformatics analysis were performed in close collaboration with the Genomics Core Facility (GCF), Norwegian University of Science and Technology (NTNU). GCF is funded by the Faculty of Medicine and Health Sciences at NTNU and Central Norway Regional Health Authority. Work at The Novo Nordisk Foundation Center for Protein Research (CPR) is funded in part by a generous donation from the Novo Nordisk Foundation (Grant number NNF14CC0001).

