## Supplementary Material for "Deciphering molecular mechanisms of synergistic growth reduction in kinase inhibitor combinations"

### Supplementary files

Supplementary Table 1: Gene Ontology terms overrepresented in PI3Ki-MEKi-specific genes at 8 hours.

| ID | Description | GeneRatio | p.adjust |
|---|---|---|---|
| GO:0042254 | ribosome biogenesis | 40/297 | 7.80E-20 |
| GO:0022613 | ribonucleoprotein complex biogenesis | 42/297 | 1.47E-18 |
| GO:0006364 | rRNA processing | 31/297 | 6.52E-16 |
| GO:0034470 | ncRNA processing | 39/297 | 1.16E-15 |
| GO:0016072 | rRNA metabolic process | 32/297 | 8.75E-15 |
| GO:0006396 | RNA processing | 46/297 | 1.01E-13 |
| GO:0034660 | ncRNA metabolic process | 48/297 | 1.11E-13 |
| GO:0042274 | ribosomal small subunit biogenesis | 12/297 | 2.40E-07 |
| GO:0000462 | maturation of SSU-rRNA from tricistronic rRNA transcript (SSU-rRNA, 5.8S rRNA, LSU-rRNA) | 9/297 | 1.78E-06 |
| GO:0030490 | maturation of SSU-rRNA | 10/297 | 1.17E-05 |
| GO:0006399 | tRNA metabolic process | 15/297 | 0.001341865831456 |
| GO:0009451 | RNA modification | 12/297 | 0.002118728218546 |
| GO:0008033 | tRNA processing | 10/297 | 0.00282203907681 |
| GO:0000460 | maturation of 5.8S rRNA | 7/297 | 0.003157097830671 |
| GO:0042273 | ribosomal large subunit biogenesis | 9/297 | 0.004408522738182 |
| GO:0006400 | tRNA modification | 8/297 | 0.012509838700026 |

Supplementary Table 2: Gene Ontology terms overrepresented in PI3Ki-TAKi-specific genes at 8 hours.

| ID | Description | GeneRatio | p.adjust |
|---|---|---|---|
| GO:0042254 | ribosome biogenesis | 37/349 | 4.67E-14 |
| GO:0022613 | ribonucleoprotein complex biogenesis | 40/349 | 6.74E-14 |
| GO:0006364 | rRNA processing | 27/349 | 8.97E-10 |
| GO:0016072 | rRNA metabolic process | 29/349 | 9.08E-10 |
| GO:0034470 | ncRNA processing | 32/349 | 1.03E-07 |
| GO:0006396 | RNA processing | 41/349 | 1.06E-07 |
| GO:0034660 | ncRNA metabolic process | 40/349 | 5.10E-06 |
| GO:0000462 | maturation of SSU-rRNA from tricistronic rRNA transcript (SSU-rRNA, 5.8S rRNA, LSU-rRNA) | 8/349 | 0.000328710111241 |
| GO:0042274 | ribosomal small subunit biogenesis | 10/349 | 0.000429838905782 |
| GO:0030490 | maturation of SSU-rRNA | 9/349 | 0.000958617354162 |
| GO:0000460 | maturation of 5.8S rRNA | 7/349 | 0.012586875267628 |
| GO:0042273 | ribosomal large subunit biogenesis | 9/349 | 0.021656053726239 |

Supplementary Figure 1: Correlation of logarithmic fold changes between each synergy and its constituting drugs, including all phosphosites that change in at least one condition.

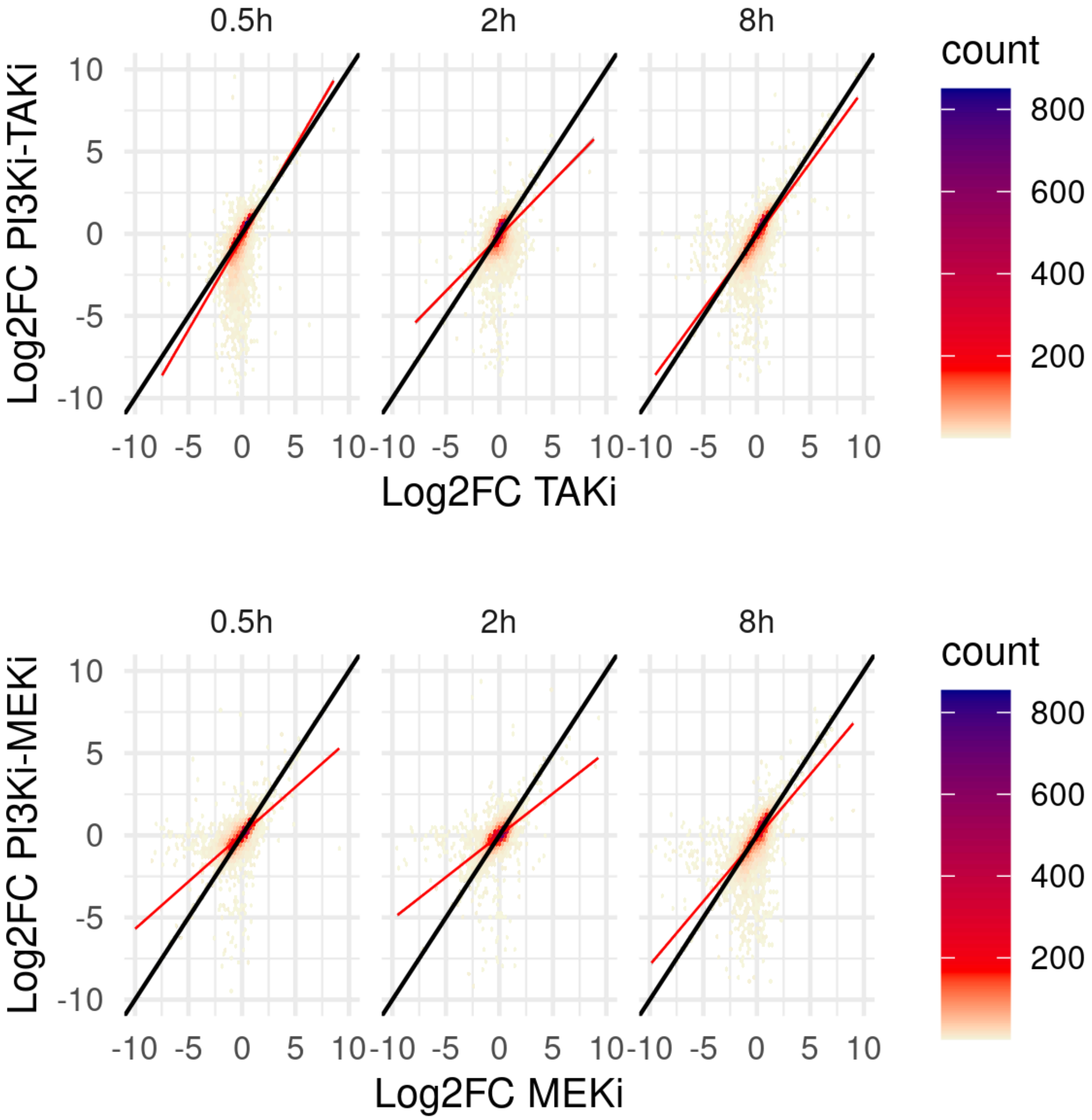

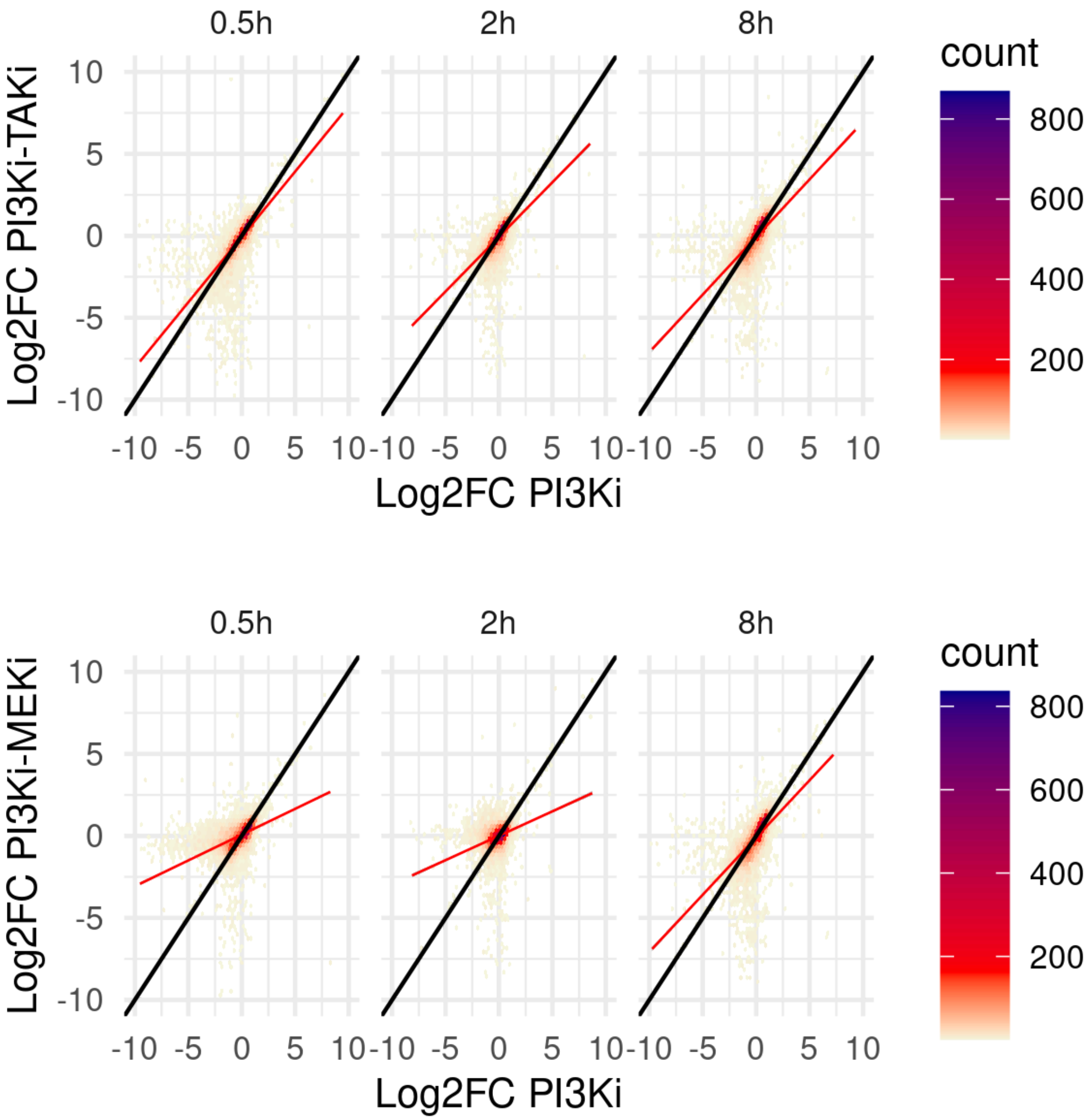
0.5h
2h
8h
count
800
600
400
200
Log2FC PI3Ki-TAKi
Log2FC PI3Ki
Log2FC PI3Ki-MEKi
10
5
0
-5
-10

Supplementary Figure 2: Schematic representation of the decision tree created to categorize changes in the synergies and the single drugs. A regulated gene refers to a gene which is differentially expressed compared to DMSO. If a gene is characterized as unaffected, the gene was not significantly upregulated or downregulated compared to DMSO.

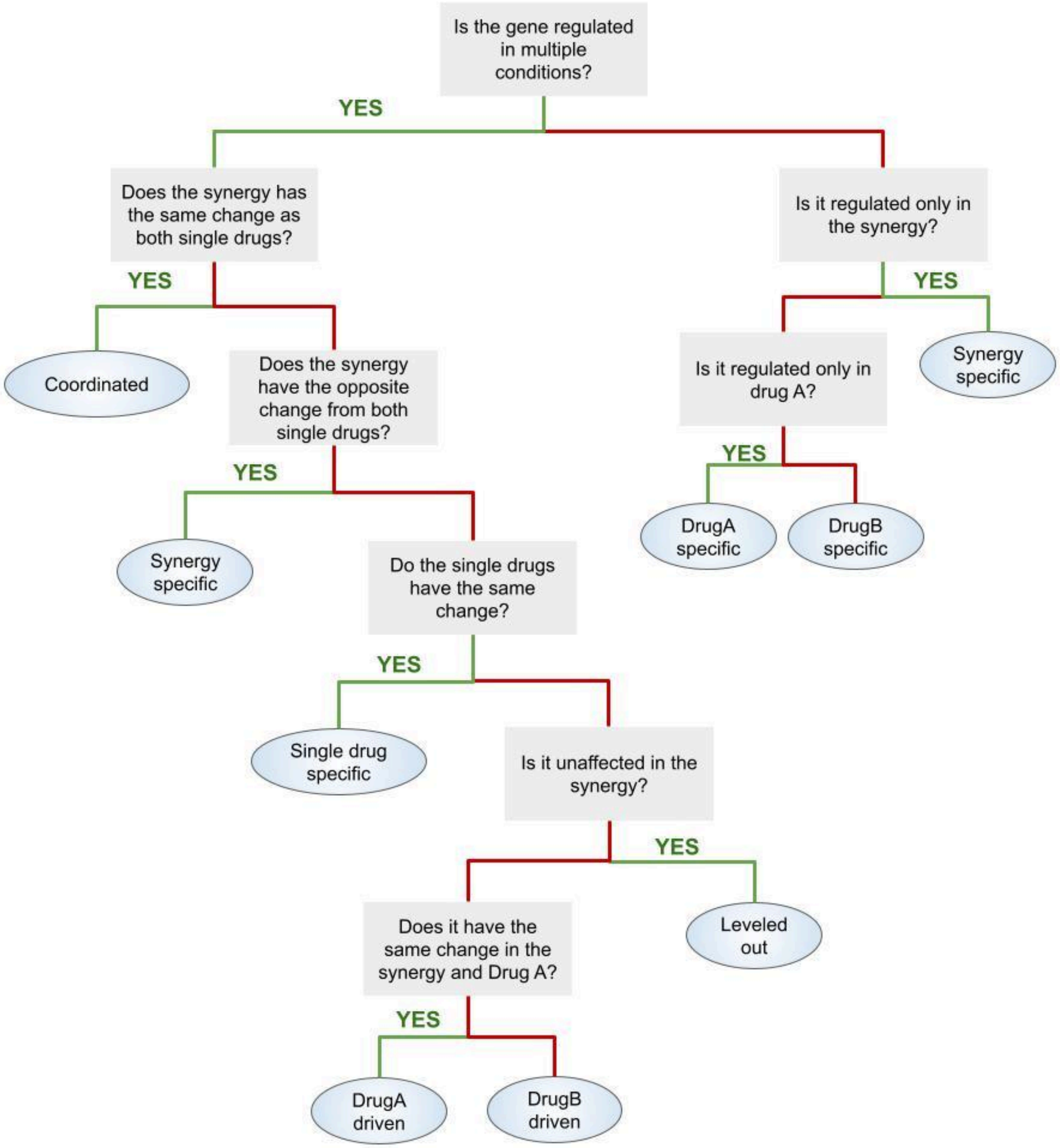

Supplementary Figure 3: Schematic representation of the response profile categories and examples for each category.

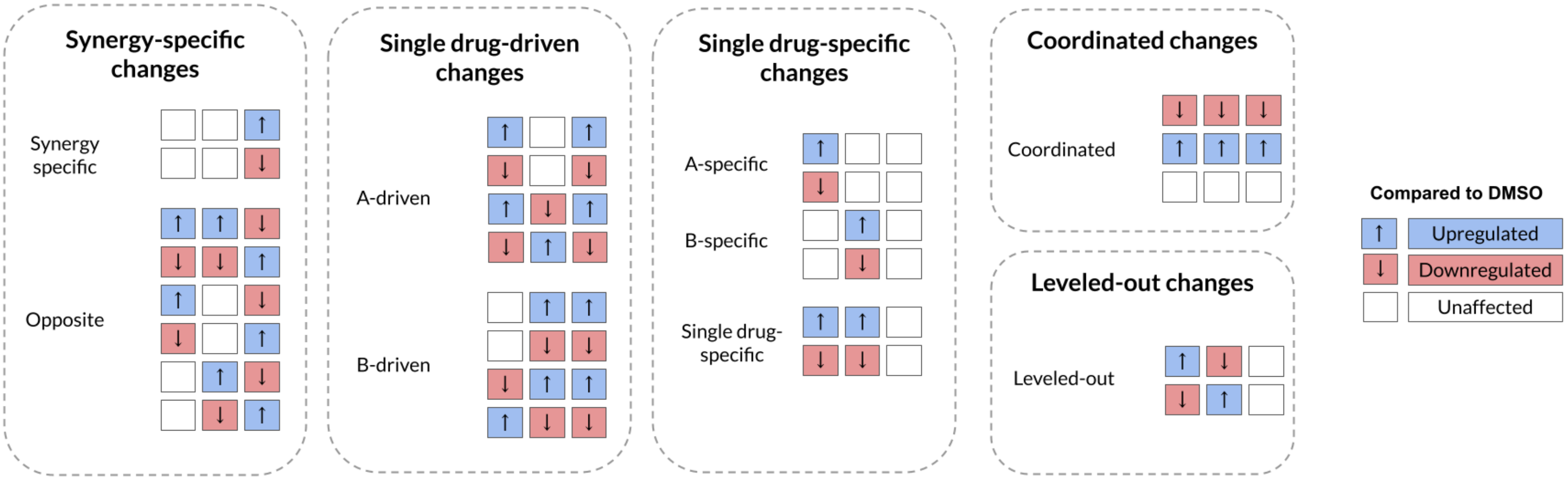

Supplementary Figure 4: Correlation of logarithmic fold changes between each synergy and its constituting drugs, including all genes that change in at least one condition.

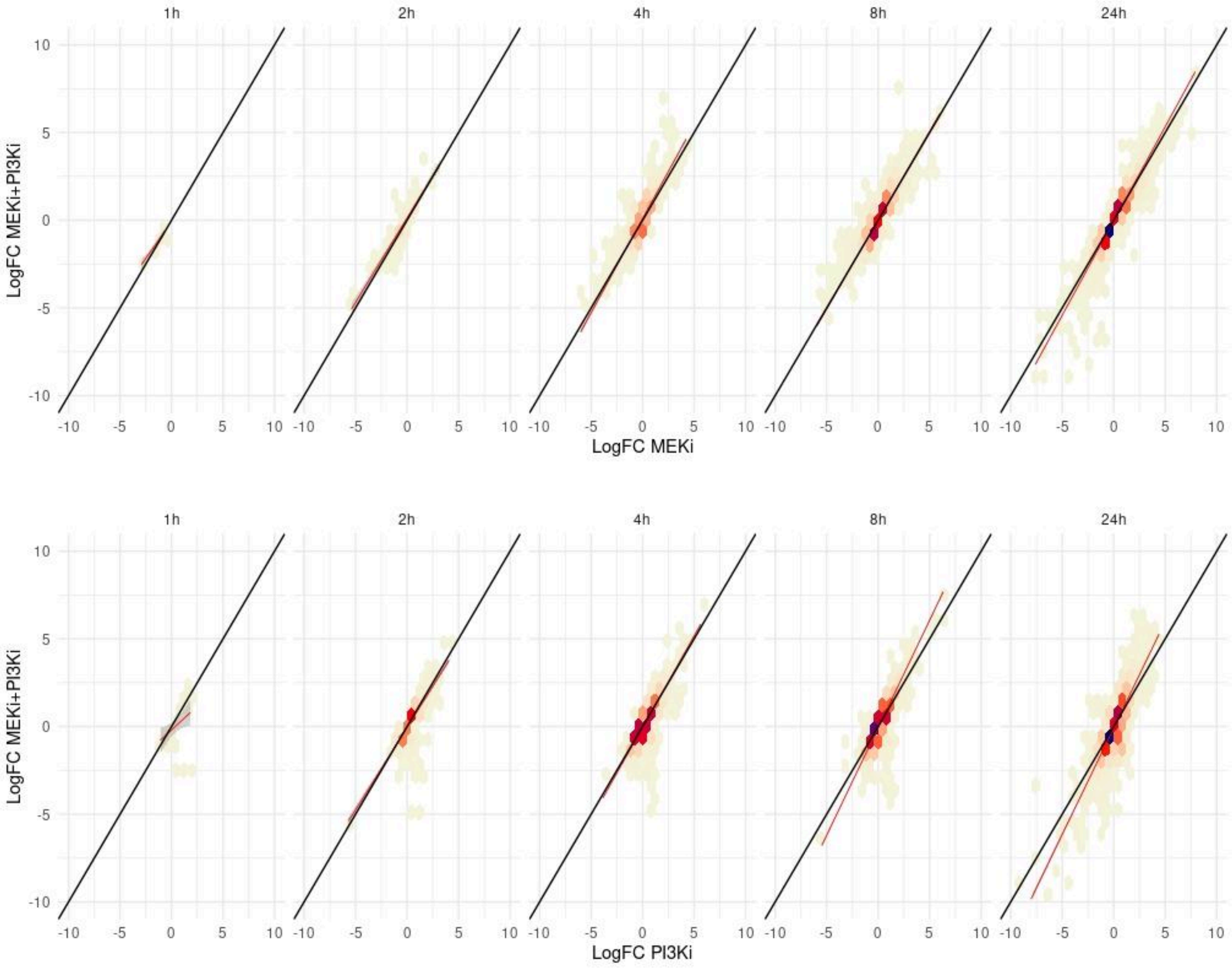

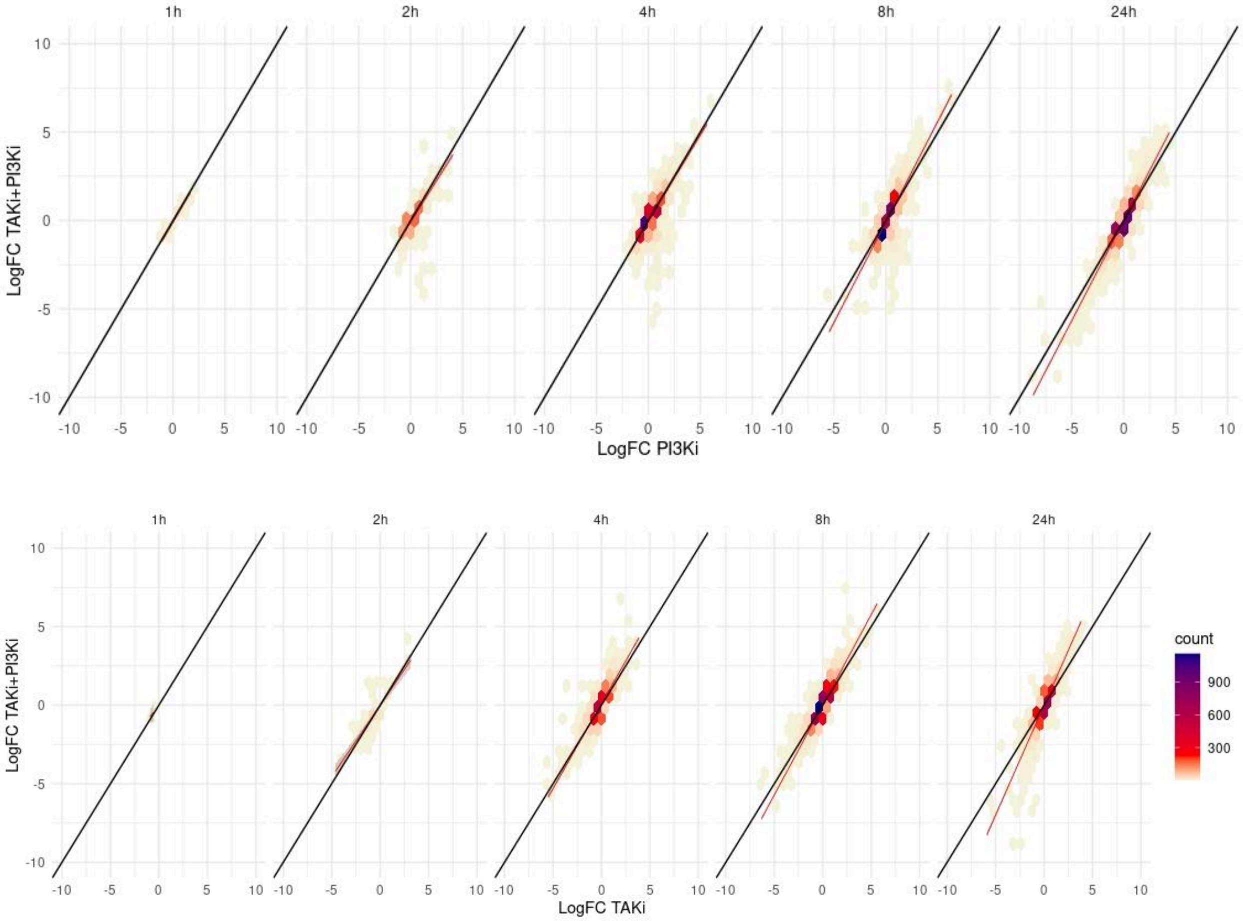
1h
2h
4h
8h
24h
LogFC TAKi+PI3Ki
LogFC PI3Ki
LogFC TAKi
count
900
600
300

Supplementary Figure 5: Estimated transcription factor activity for all conditions

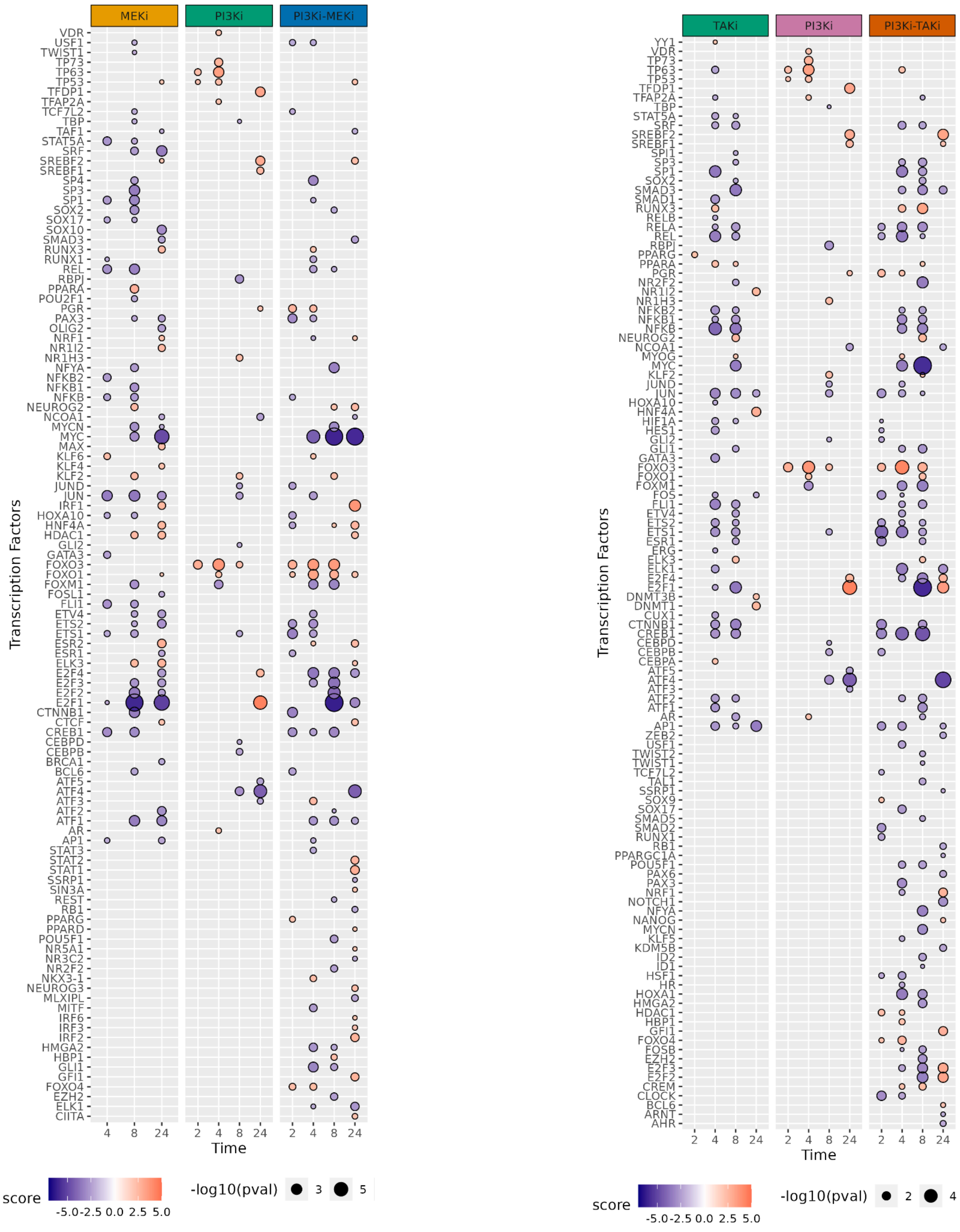

Supplementary Figure 6: Logarithmic fold change (logFC) of genes involved in nucleotide metabolism. LogFCs were calculated with DMSO as a baseline.

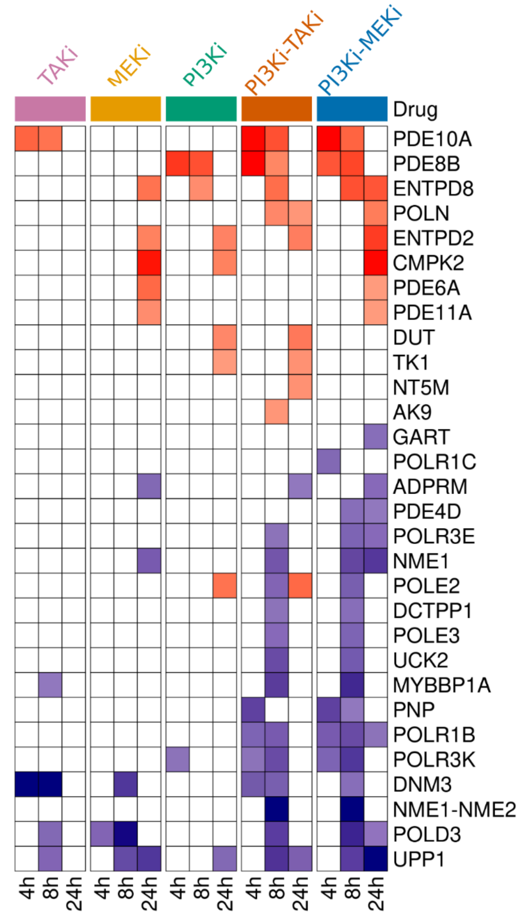

Supplementary Figure 7: Heatmap of the logarithmic fold changes of gene expression when compared to DMSO, for genes annotated with autophagy-related Gene Ontology terms.

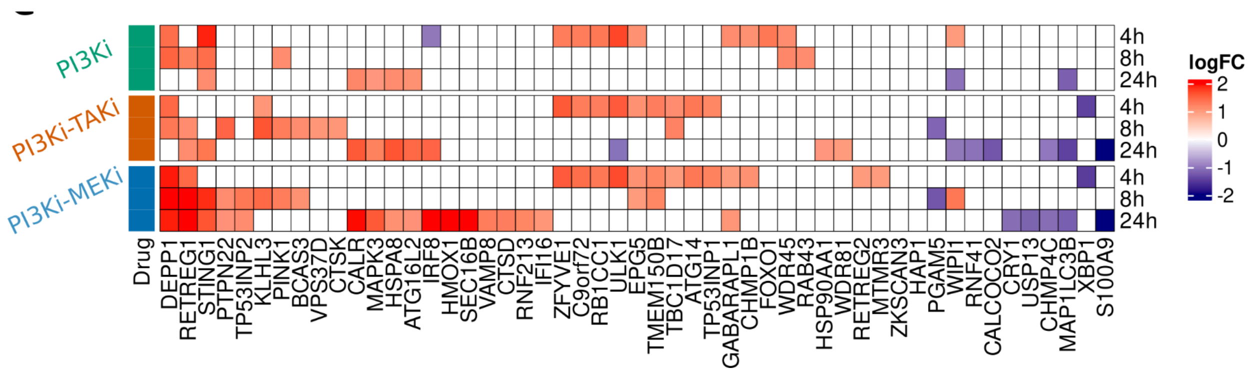
